# Localized inhibition in the *Drosophila* mushroom body

**DOI:** 10.1101/2020.03.26.008300

**Authors:** Hoger Amin, Raquel Suárez-Grimalt, Eleftheria Vrontou, Andrew C. Lin

## Abstract

Many neurons show compartmentalized activity, in which activity does not spread readily across the cell, allowing input and output to occur locally. However, the functional implications of compartmentalized activity for the wider neural circuit are often unclear. We addressed this problem in the *Drosophila* mushroom body, whose principal neurons, Kenyon cells, receive feedback inhibition from a large, non-spiking interneuron called APL. We used local stimulation and volumetric calcium imaging to show that APL inhibits Kenyon cells in both their dendrites and axons, and that both activity in APL and APL’s inhibitory effect on Kenyon cells are spatially localized, allowing APL to differentially inhibit different mushroom body compartments. Applying these results to the *Drosophila* hemibrain connectome predicts that individual Kenyon cells inhibit themselves via APL more strongly than they inhibit other individual Kenyon cells. These findings reveal how cellular physiology and detailed network anatomy can combine to influence circuit function.

## Introduction

A textbook neuron integrates input from dendrites to generate action potentials that spread throughout the neuron, making it a single functional unit. However, many neurons integrate inputs and generate outputs locally so that a single neuron effectively functions as multiple independent units (reviewed in (Branco and Häusser, 2010)). What functional consequences flow from such compartmentalized activity? In some cases, the circuit function of compartmentalized activity is clear (Euler et al., 2002; Grimes et al., 2010), but answering this question is often difficult due to the complexity of mammalian nervous systems or lack of detailed anatomical information.

These difficulties are eased in the *Drosophila* olfactory system, a numerically simple circuit that is now subject to intensive connectomic reconstruction (Takemura et al., 2017; Tobin et al., 2017; Xu et al., 2020; Zhang et al., 2019; Zheng et al., 2018), offering an excellent opportunity to study local computations. We focus on the mushroom body, the fly’s olfactory learning center, which has a well-characterized compartmentalized architecture (Amin and Lin, 2019). The mushroom body’s principal neurons, the Kenyon cells (KCs), form parallel bundles of axons that are divided into compartments by the innervation patterns of dopaminergic neurons and mushroom body output neurons. During learning, dopaminergic neurons in one compartment modify the local connection strength between Kenyon cells and mushroom body output neurons in the same compartment, but not other compartments (Cohn et al., 2015; Hige et al., 2015).

Superimposed on this compartmentalized structure is an inhibitory interneuron that innervates all the mushroom body compartments: the anterior paired lateral neuron (APL) (Aso et al., 2014; Tanaka et al., 2008). The APL neuron ensures sparse odor coding by Kenyon cells through feedback inhibition (Lei et al., 2013; Lin et al., 2014) and also plays a role in memory suppression and reversal learning (Liu and Davis, 2008; Ren et al., 2012; Wu et al., 2012; Zhou et al., 2019). APL’s widespread innervation stands in contrast to the generally compartmentalized structure of the mushroom body, yet its diverse roles in memory hint that it may have a more compartmentalized function than its morphology suggests.

Indeed, there are hints that APL may have compartmentalized function. Like its locust homolog, the giant GABAergic neuron (GGN), APL is non-spiking (Papadopoulou et al., 2011). Unlike the larval APL (Eichler et al., 2017; Masuda-Nakagawa et al., 2014), the adult APL has pre- and post-synaptic specializations throughout all its processes (Wu et al., 2013), and connectomic reconstructions show that APL forms reciprocal synapses with Kenyon cells throughout the mushroom body (Takemura et al., 2017; Xu et al., 2020; Zheng et al., 2018). There have been reports of localized activity within APL (Inada et al., 2017; Wang et al., 2019), and compartmental modelling suggests that in the locust GGN, depolarization should diminish with distance from the site of current input, though not to zero (Ray et al., 2019). However, intracellular activity propagation is difficult to measure experimentally in locusts, and the lack of action potentials in APL at the soma (Papadopoulou et al., 2011) does not rule out active conductances elsewhere, given that mammalian dendrites often show dendritic spikes that do not necessarily propagate to the soma (Branco and Häusser, 2010). There have been no systematic studies of how widely activity propagates within APL, where it exerts its inhibitory effect on Kenyon cells, or how APL’s physiology would affect the structure of feedback inhibition in the mushroom body.

We addressed these questions by volumetric calcium imaging in APL. By quantifying the spatial attenuation of the effects of locally stimulating APL, we found that both activity in APL and APL’s inhibitory effect on Kenyon cells are spatially restricted, allowing APL to differentially inhibit different compartments of the mushroom body. Combining these physiological findings with recent connectomic data predicts that individual Kenyon cells inhibit themselves via APL more strongly than they do other Kenyon cells. Our findings establish APL as a model system for studying local computations and their role in learning and memory.

## Results

### Spatially restricted responses to electric shock in APL

We first asked whether physiological responses to sensory stimuli are spatially localized within APL. At very low odor concentrations, odor responses in APL can be restricted to one lobe (Inada et al., 2017), but it remains unclear how sensory-evoked activity is structured across APL at larger scales. We examined this question using electric shock, a typical ‘punishment’ used for olfactory aversive conditioning.

APL responds to electric shock (Liu and Davis, 2008; Zhou et al., 2019), but it is unknown if these responses are spatially restricted. To test this, we subjected flies to electric shock and recorded activity volumetrically throughout the APL lobes using GCaMP6f driven by 474-GAL4 (Gohl et al., 2011). We divided the MB lobes into 4 regions: the tip of the vertical lobe (V), stalk of the vertical lobe (S), compartments γ1-3 (γ), and the rest of the horizontal lobe (H) (**Fig. 1A**). We based these divisions on findings that (1) these regions are innervated by different dopaminergic neurons that respond differentially to ‘punishment’ (e.g., electric shock) vs. ‘reward’ (reviewed in (Amin and Lin, 2019)), (2) dopaminergic neurons release synaptic vesicles onto APL in response to electric shocks (Zhou et al., 2019), and (3) APL expresses the dopamine receptors DopEcR (Aso et al., 2019) and the inhibitory Dop2R (Zhou et al., 2019).

**Figure 1.**
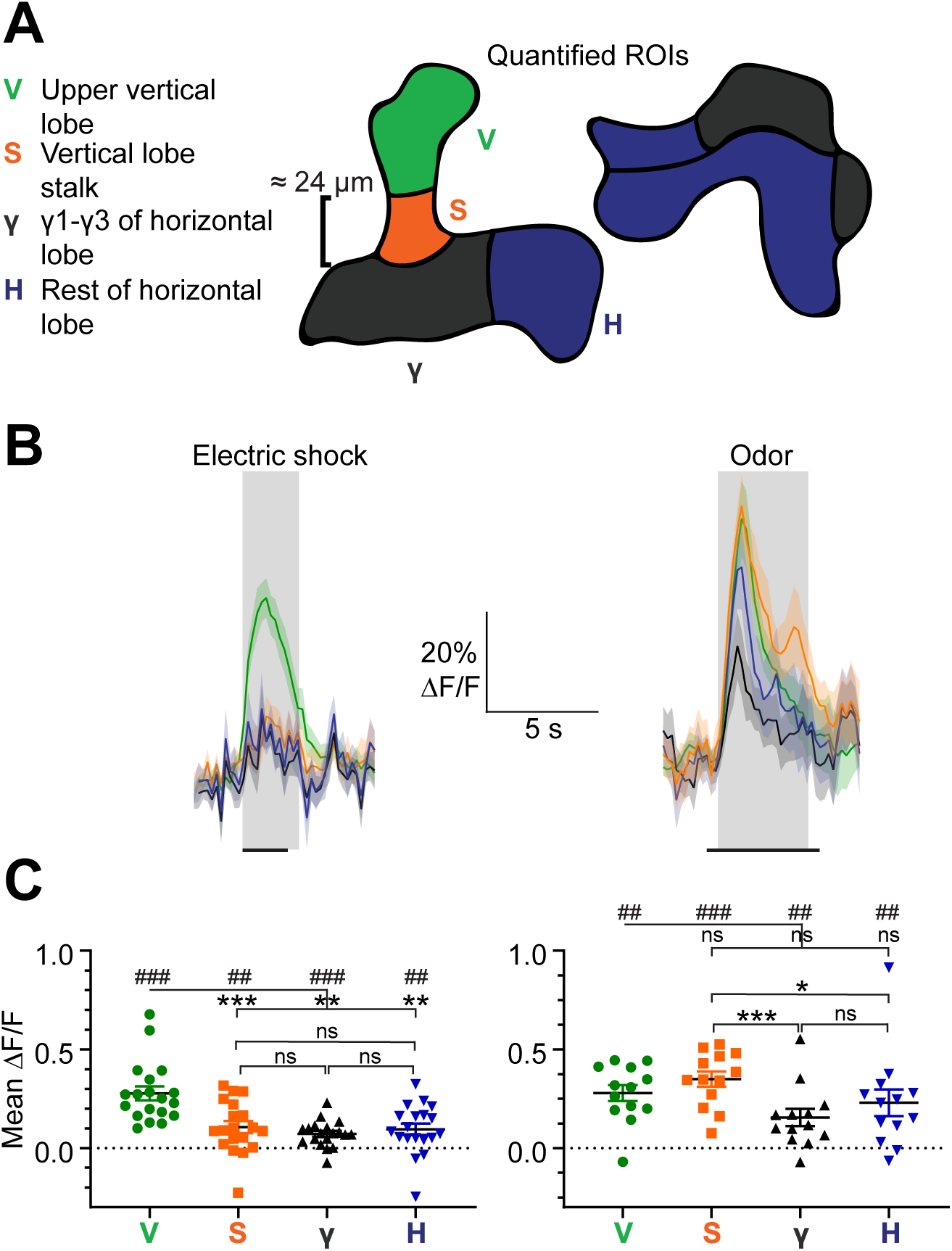
Spatially restricted responses to electric shock in APL. **(A)** Diagrams of regions used to quantify APL activity in **(B-C)**. **(B)** APL responses to electric shock (left) or the odor isoamyl acetate (right) in the regions defined in (A), in 474-GAL4>GCaMP6f flies. Traces show mean ± SEM shading. Electric shock trace is the average of 3 shocks with 12 s between onset of each shock. Horizontal black lines show timing of electric shock (1.2 s) or odor (5 s). **(C)** Mean ΔF/F during the intervals shaded in panel **(B)**. ## p<0.01, ### p<0.001, one-sample Wilcoxon test vs null hypothesis (0) with Holm-Bonferroni correction for multiple comparisons. * p<0.05, *** p<0.001, Friedman t-test with Dunn’s multiple comparisons test for each group (V, H, S, and γ) against each other. n, given as # neurons (# flies): electric shock, 19 (15); odor, 13 (11).

Electric shocks evoked a strong response in APL in the V region, with significantly smaller responses in the other regions (**Fig. 1B,C**). This difference between regions did not arise from optical artifacts or differential expression of voltage-gated calcium channels. If it did, one would expect other stimuli to produce a similar pattern of Ca^2+^ influx. However, the odor isoamyl acetate evoked a different pattern of activity, with somewhat stronger responses in the S region and weaker in γ **(Fig. 1B,C)**. These different spatial profiles for electric shock v. odor suggest that spatially differential responses to electric shock might arise from different synaptic inputs. This could arise because different regions of APL (1) have dopamine receptors of different function (DopEcR vs. Dop2R), (2) receive dopaminergic inputs of different synaptic numbers, or (3) receive input from dopaminergic neurons with different responses to electric shock (Cohn et al., 2015; Mao and Davis, 2009). In support of (1), bath-applying dopamine inhibits APL in the vertical lobe, except in the tip where it has no direct effect (Zhou et al., 2019). In support of (2), APL receives more synapses from the dopaminergic neurons PPL1γ2α′1, PPL1γ1pedc and PPL1α′2α2 in the heel and stalk, which are presumably inhibitory (Zhou et al., 2019), compared to dopaminergic neurons PPL1α3 and PPL1α′3 in the tip (Xu et al., 2020)). Regardless of the cause, the spatial heterogeneity of sensory-evoked activity within APL suggests that activity can remain spatially localized in APL.

### Low expression of voltage-gated Na^+^ and Ca^2+^ channels in APL

In addition to reflecting differential synaptic inputs, this spatially localized activity in APL most likely also reflects intrinsic properties of APL’s membrane excitability that prevent activity from spreading. To validate the hypothesis that intrinsic properties could contribute, we analyzed an RNA-seq dataset revealing gene expression in Kenyon cells, dopaminergic neurons, mushroom body output neurons, the dorsal paired medial (DPM) neuron, and APL (Aso et al., 2019).

We looked for genes with higher or lower expression in APL than in all other 20 mushroom body neuron types. Gene Ontology analysis revealed that the set of genes with consistently higher expression in APL is strongly enriched for genes involved in protein synthesis and mitochondrial respiration, perhaps reflecting the energetic demands of maintaining such a large neuron. APL also expresses higher levels of the biosynthetic enzyme and vesicular transporter of GABA than any other neuron in the dataset (**Fig. 2**). In contrast, voltage-gated Na^+^ and Ca^2+^ channels are expressed at lower levels in APL than in all other types of mushroom body neurons (**Fig. 2**), indeed 10-1000x lower than the average of non-APL neurons. In contrast, the picture with K^+^ channels, ligand-gated Cl^-^ channels, and other miscellaneous channels is more mixed (**Fig. 2**). These results are consistent with previous findings that APL does not fire action potentials (Papadopoulou et al., 2011), unlike all other published electrophysiological recordings from other mushroom body neurons (Hige et al., 2015; Takemura et al., 2017; Turner et al., 2008).

**Figure 2.**
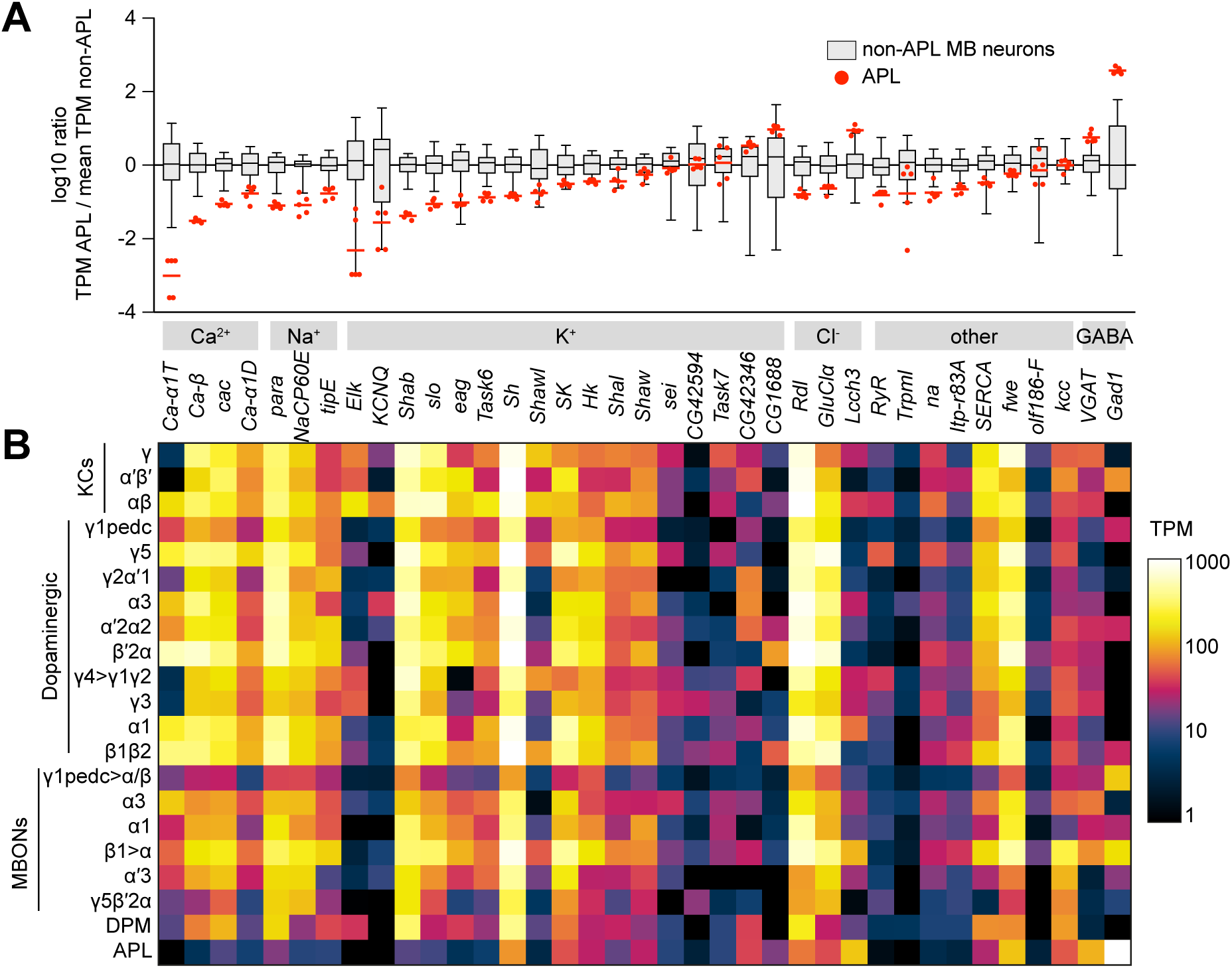
Low expression of voltage-gated Na^+^ and Ca^2+^ channels in APL. **(A)** Expression levels of GABAergic markers and various Na^+^, Ca^2+^, K^+^, Cl^-^ and other channels, in APL compared to non-APL mushroom body neurons, normalized to the mean of non-APL neurons. Box plots show the distribution of expression levels in non-APL MB neurons (whiskers show full range), using the average of log_10_(TPM) across biological replicates for each type. Red dots show 5 biological replicates of APL; red line shows the mean APL expression level. Genes are grouped by type and sorted by fold difference in APL expression relative to non-APL MB neurons. Data from (Aso et al., 2019). TPM, transcripts per million. **(B)** Heat map of TPM counts (log scale) for all genes shown in (A), for all 21 cell types.

### Activity in APL is spatially restricted

Based on the spatially restricted sensory-evoked activity and the low expression of voltage-gated ion channels, we hypothesized that intrinsic factors limit the spatial spread of activity within APL. We first tested this by imaging APL while activating Kenyon cells with the heat-activated cation channel dTRPA1. We previously showed that activating all Kenyon cells with dTRPA1 activates APL (Lin et al., 2014). We now extended this approach by activating only subsets of Kenyon cells. Kenyon cells are divided into three main classes, αβ, α′β′, and γ, which send their axons to anatomically segregated mushroom body lobes, called the α, β, α′, β′, and γ lobes. When we activated only the α′β′ Kenyon cells and imaged APL in the vertical lobe (made of the α and α′ lobes), remarkably, only the α′ lobe of APL responded (**Fig. 3A-C,I**). The same occurred for the α lobe when we activated only αβ/γ (**Fig. 3D-F,I**) or only αβ Kenyon cells (**Fig. 3G-I**). APL neurites in the unresponsive lobe failed to respond despite being only 2-3 µm away from responding neurites. This can most likely be explained by the fact that APL neurites in the vertical lobe mostly run in parallel along Kenyon cells and do not cross between the α and α′ lobes (Takemura et al., 2017). These results suggest that activity in APL fails to propagate over long distances.

**Figure 3.**
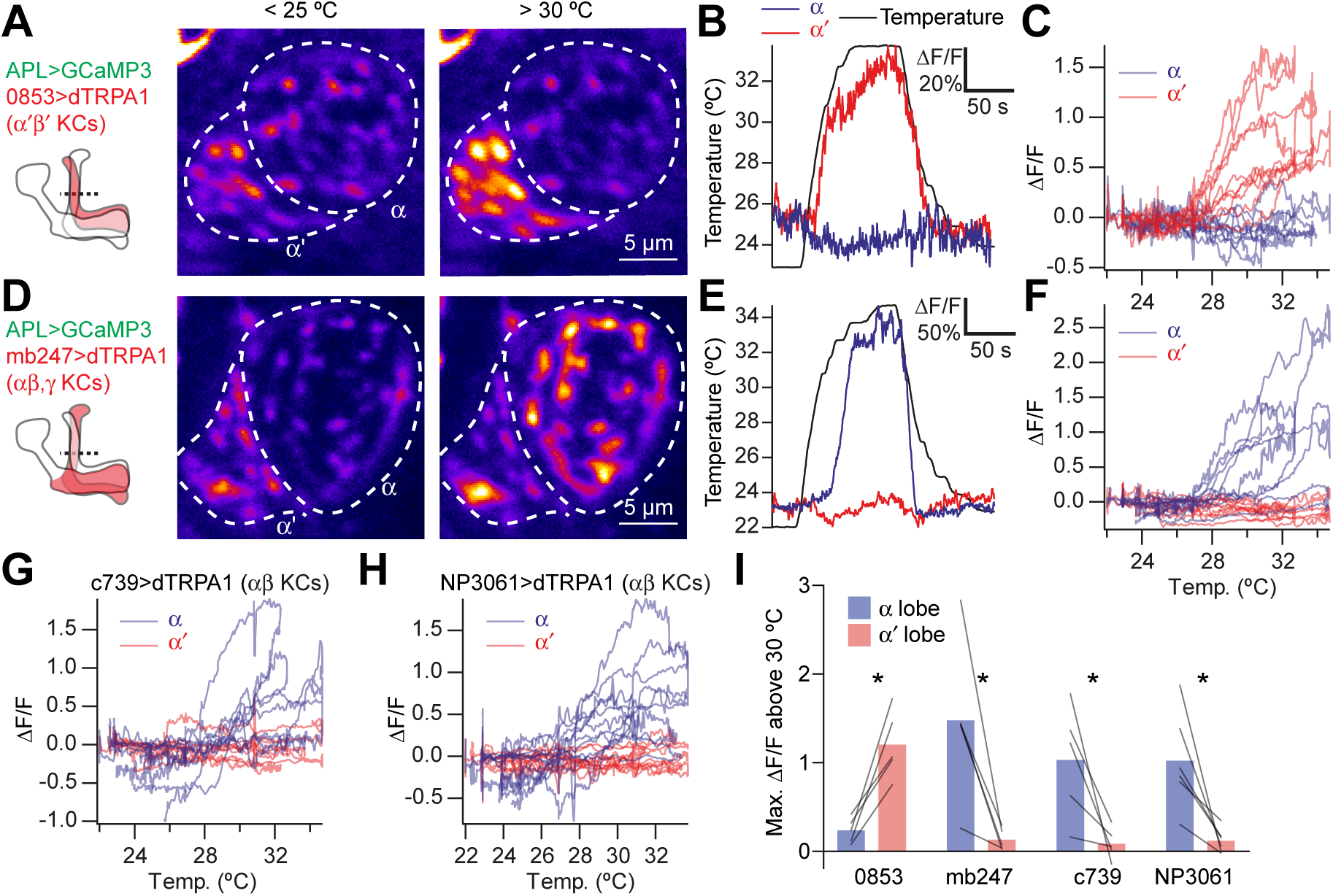
Activating subsets of Kenyon cells only activates the corresponding lobe of APL. **(A)** Time-averaged images of APL in a cross-section of the lower vertical lobe (dotted line in diagram) at room temperature (< 25 °C) vs. heated (> 30 °C). Activating α′β′ Kenyon cells expressing 0853-GAL4-driven dTRPA1 activates the α′ lobe, but not the α lobe, of APL. APL expresses GCaMP3 driven by GH146-QF. **(B)** Example responses to heating (black), by the α′ (red) and α (blue) lobes in the same APL recording. **(C)** ΔF/F for the α′ (red) and α (blue) lobes plotted against temperature. Each trace represents one recording forms a loop: it rises as the fly is heated and falls down a different path as the fly is cooled. **(D-F)** As for **(A-C)** except activating αβ and γ Kenyon cells (mb247-GAL4>dTR-PA1) drives activity in the α, but not the α′ lobe of APL. **(G,H)** As for **(C)** and **(F)** except activating αβ Kenyon cells with c739-GAL4 (G) or NP3061-GAL4 (H). **(I)** Maximum ΔF/F between 30 °C and the peak temperature, summarizing data from **(C,F-H)**. Lines indicate the same APL recording. * p < 0.05, paired t-test with Holm-Bonferroni multiple comparisons correction. n (# flies), left to right: 5, 5, 5, 6.

To measure activity propagation within APL more systematically, we directly stimulated APL, to test whether local activation of APL in the mushroom body lobes (where Kenyon cell axons reside) would spread to the calyx (where Kenyon cell dendrites reside), and vice versa (**Fig. 4**). To do this, we expressed GCaMP6f (Chen et al., 2013) and P2×2, an ATP-gated cation channel (Lima and Miesenböck, 2005), in APL, using a previously described intersectional driver for specifically labeling APL (NP2631-GAL4, GH146-FLP, tubP-FRT-GAL80-FRT (Lin et al., 2014)). *Drosophila* does not natively express P2X ATP receptors (Littleton and Ganetzky, 2000). We locally applied ATP to the tip of the vertical lobe or to the calyx by pressure ejection from a Picospritzer, while imaging both regions. This local stimulation evoked intense GCaMP6f responses at the site of ATP application, but not at the unstimulated site **(Fig. 4A,D)**, indicating that activity in APL attenuates to undetectable levels across the breadth of the neuron (about 250-300 µm; see below).

**Figure 4.**
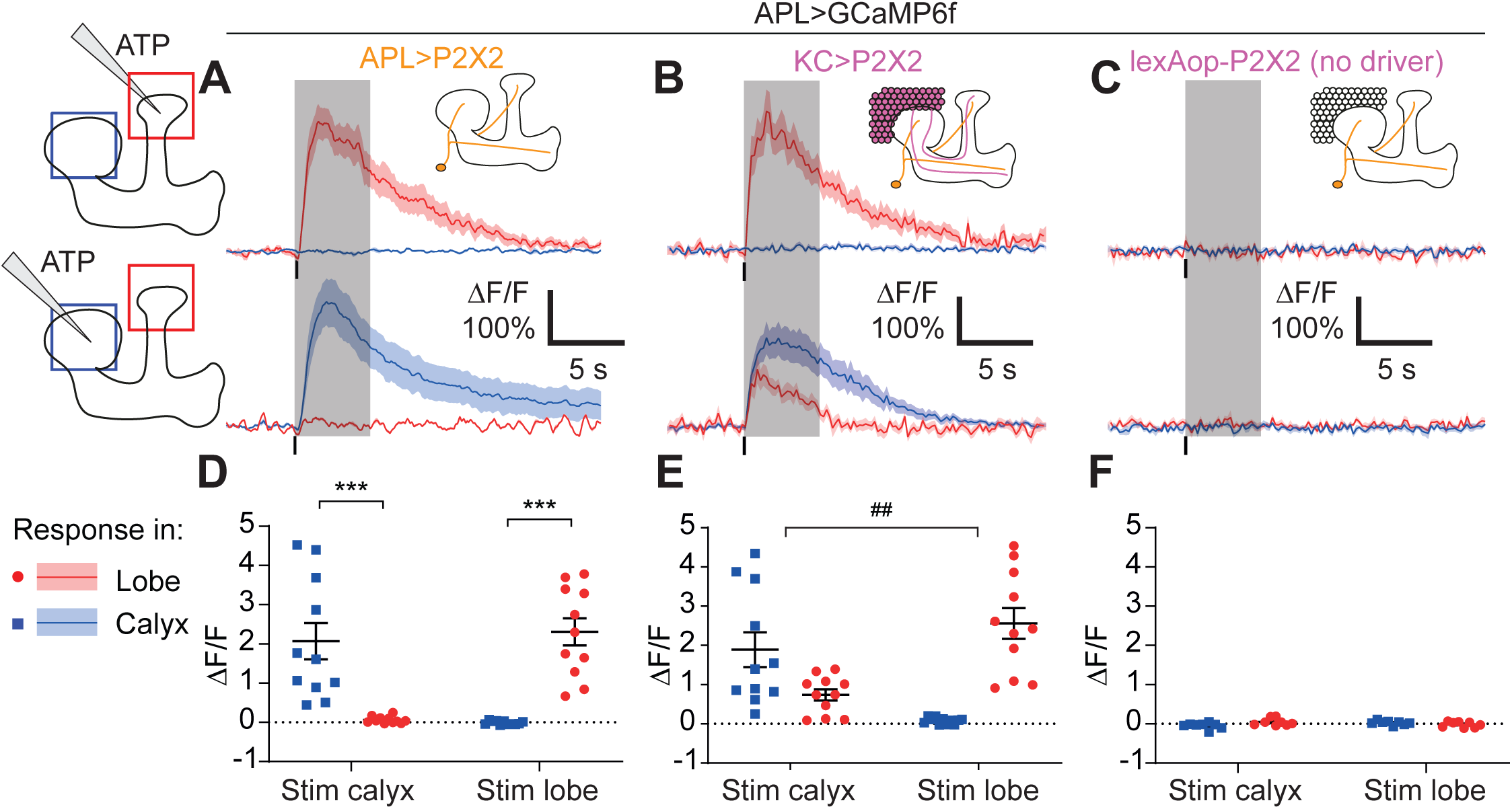
Activity in the APL neuron is spatially restricted. **(A-C)** Response of APL in the vertical lobe (red) or calyx (blue) to pressure ejection of 1.5 mM ATP (100 ms pulse, 12.5 psi) in the lobe or calyx, to stimulate APL **(A)**, KCs **(B)**, or negative control flies with lexAop-P2×2 but no LexA driver **(C)**. Schematics show ROIs where responses were quantified (red/blue squares), and where ATP was applied (pipette symbol). Traces show mean ± SEM shading. Vertical lines on traces show time of ATP application; gray shading shows 5 s used to quantify response in **(D-F)**. **(D-F)** Mean ΔF/F from **(A-C)**, when stimulating APL **(D)**, KCs **(E)**, or negative controls **(F)**. Error bars show SEM. n, given as # neurons (# flies): **(A,D)**: 11 (9); **(B,E)**: 11 (7); **(C,F)** 8 (4). *** p<0.001, t-tests with Holm Sidak multiple comparisons correction. ## p = 0.004, paired t-test: the ratio of (response in unstimulated site)/(response in stimulated) site is higher for calyx stimulation than lobe stimulation.

To rule out the possibility that all P2×2-driven excitation is always spatially restricted, we repeated the experiment but expressed P2×2 in Kenyon cells instead of APL, using mb247-LexA (Pitman et al., 2011). We still expressed GCaMP6f in APL, now using 474-GAL4 instead of the intersectional driver, to increase throughput (as this experiment did not require highly specific expression in APL). In contrast to stimulating APL, stimulating Kenyon cells in the calyx evoked a strong response in APL in both the calyx and the vertical lobe (**Fig. 4B,E)**, most likely because activating Kenyon cells in their dendrites drives them to spike, which then activates APL throughout the mushroom body. However, stimulating Kenyon cells at the tip of the vertical lobe activated APL locally, but not in the calyx (**Fig. 4B,E)**. Puffing ATP on negative control flies with no driver for P2×2 evoked no responses in APL (**Fig. 4C,F**). Together, these findings show that activity within APL is spatially restricted.

### Quantification of activity spread in APL

Locally induced activity in APL declines to essentially zero at the other end of the neuron, but how quickly does the activity decay along its journey? In particular, is activity in APL restricted enough for APL to signal differentially in different compartments of the MB lobes? We examined this question by more systematically quantifying spread of activity throughout APL. We used volumetric imaging to record activity in the entire 3D extent of APL (∼100 x 100 x 150 µm). We analyzed APL as a 3D ‘skeleton’ with three branches (vertical lobe, horizontal lobe, and peduncle/calyx). We used evenly spaced points every 20 µm along this skeleton to divide the 3D volume of APL into segments (Voronoi cells), and quantified fluorescence signals in APL in each segment-volume (**Fig. 5A**, see Methods). To compare the spatial extent of activity across flies, we normalized distances in these skeletons to a ‘standard’ skeleton based on the average across all flies (**Fig. 5B**). Most neurites in APL run in parallel with this skeleton (Takemura et al., 2017; Wang et al., 2019; Xu et al., 2020), making this skeleton a meaningful axis along which to measure spread of APL activity.

**Figure 5.**
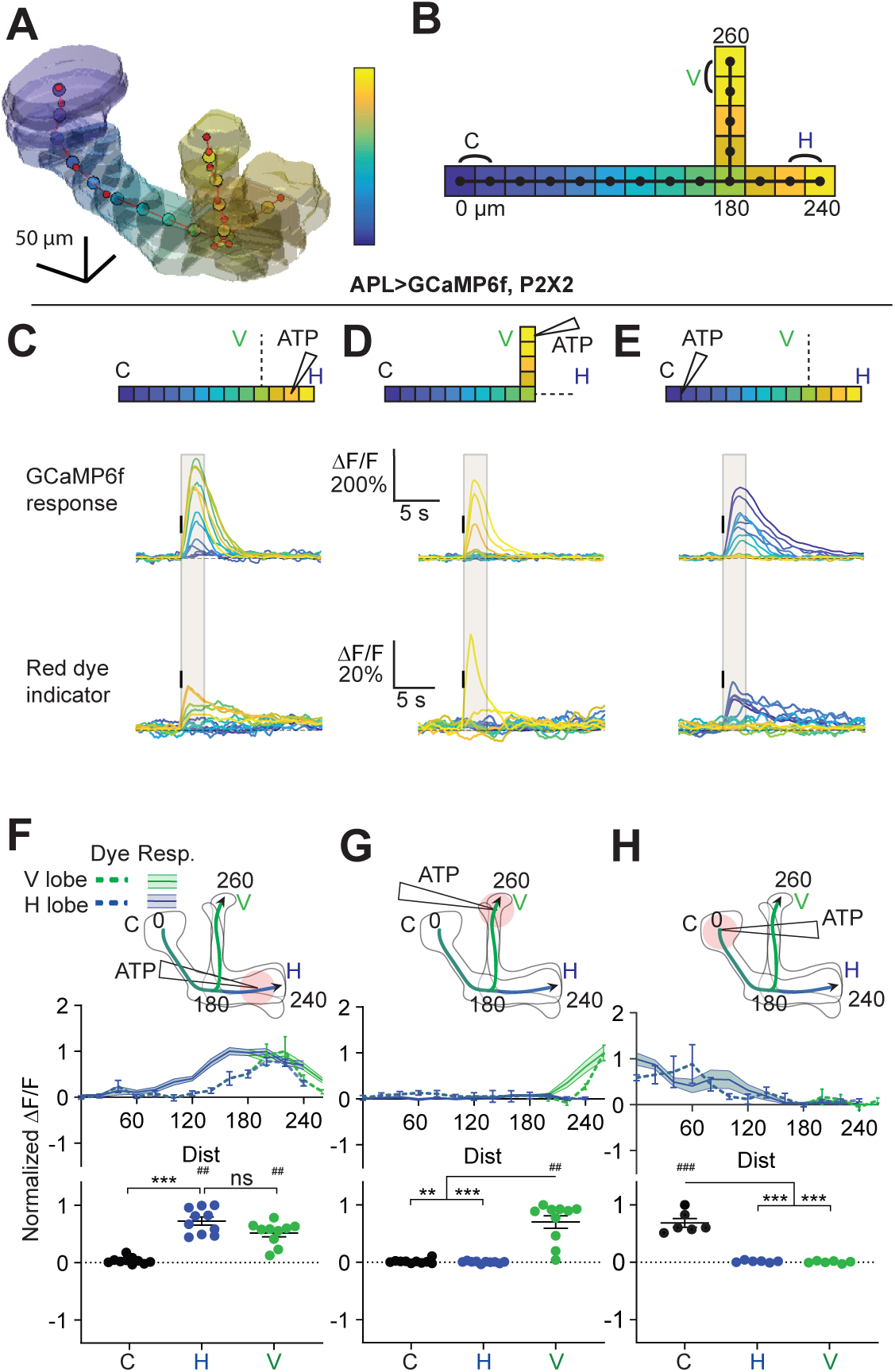
Quantification of activity spread in APL. **(A)** 3D visualization of an example mushroom body manually outlined using mb247-dsRed as an anatomical landmark. The skeleton is shown as a red line passing through the center of each branch of the mushroom body, manually defined by the red dots. Evenly spaced nodes every 20 µm on the skeleton subdivide the mushroom body into segments, here color-coded according to distance from the dorsal calyx (scale on right). **(B)** Schematic of the ‘standard’ 3D skeleton with distances (µm) measured from the dorsal calyx. Color code as in panel A. In this and later figures, signals are quantified using the two outermost segments of the calyx (C, black), vertical lobe (V, green), and horizontal lobe (H, blue), as shown here. **(C-E)** Traces show the time course of the response in each segment of APL, averaged across flies, when stimulating APL with 0.75 mM ATP (10 ms puff at 12.5 psi) in the horizontal lobe **(C)**, vertical lobe **(D)**, or calyx **(E)**. Upper panels: GCaMP6f signal. Lower panels: Red dye signal. The baseline fluorescence for the red dye signal comes from mb247-dsRed. Color-coded skeleton indicates which segments have traces shown (dotted lines mean the data is omitted for clarity; the omitted data appear in panels **F-H**). Gray shading shows the time period used to quantify ΔF/F in **(F-H)**. Vertical black lines indicate the timing of ATP application. **(F-H)** Mean response in each segment (ΔF/F averaged over time in the gray shaded period in **C-E**). The color of the curves matches the vertical (green) and horizontal (blue) lobular branches depicted in the schematics. The responses in each panel were normalized to the highest responding segment (upper panels) or data point (lower panels). Error bars show SEM. n, given as # neurons (# flies): **(C, F, D, G)** 10 (6), **(E, H)** 6 (4). ## p<0.01, ### p<0.001, one-sample Wilcoxon test or one-sample t-test, vs null hypothesis (0) with Holm-Bonferroni correction for multiple comparisons. ** p<0.01, *** p<0.001, Friedman t-test with Dunn’s multiple comparisons test or repeated measures one-way ANOVA with Holm-Sidak’s multiple comparisons test, comparing the stimulated vs the unstimulated sites. See Table S2 for detailed statistics.

To avoid the stochastic labeling of APL associated with the intersectional NP2631/GH146 driver, we developed another APL driver, VT43924-GAL4.2-SV40 (henceforth simply VT43924-GAL4.2), based on the original VT43924-GAL4 (Wu et al., 2013), as GAL4.2-SV40 typically gives ∼2x stronger expression than the original GAL4 (Pfeiffer et al., 2010). This line showed more reliable expression than the original VT43924-GAL4 line and labelled fewer neurons near the MB compared to 474-GAL4 (**Figure S1A,B**), making it applicable for ATP stimulation of APL. We used VT43924-GAL4.2 for labeling APL in all subsequent experiments.

We locally activated APL by applying ATP to three zones – the tip of the vertical lobe, the calyx, and the horizontal lobe – and quantified the response at evenly spaced intervals along the 3D skeleton. ATP stimulation of APL evoked a strong local response that gradually decayed as it propagated away from the stimulus site (**Fig. 5**), with little change in the temporal dynamics of the response with distance (**Fig. 5C-E**). Note that our temporal resolution (∼5 Hz imaging rate) was not fast enough to reveal propagation of activity along APL neurites over time (which presumably occurs on the timescale of milliseconds). We co-ejected a red dye together with the ATP to reveal the spatial and temporal extent of stimulation as the ATP was removed by diffusion and perfusion. For example, the longer response from stimulation in the calyx reflects the slower disappearance of ATP (**Fig. 5C-E**, lower panels).

Strikingly, activity evoked by stimulation at the tip of the vertical lobe decayed to an undetectable level by the branching point between the two lobes (**Fig. 5G**). Stimulation in the other two zones elicited wider spread, but much of the activity spread actually arises from movement of ATP away from the ejection site, as revealed by spread of the co-ejected red dye (dashed lines, **Fig. 5**). These results confirm the spatially restricted activity shown in **Fig. 4** and extend those results by systematically quantifying the extent of activity spread throughout APL.

### APL inhibits Kenyon cells mostly locally

Given that activity within APL is localized, we next asked whether APL’s inhibitory effect on Kenyon cells is also localized. APL and Kenyon cells form reciprocal synapses throughout the entire extent of the mushroom body neuropil (**Fig. 8**) (Takemura et al., 2017; Xu et al., 2020; Zheng et al., 2018), suggesting that APL could theoretically inhibit Kenyon cells everywhere. However, it remains unknown if APL actually can inhibit Kenyon cells everywhere and if this inhibition remains local.

**Figure 6.**
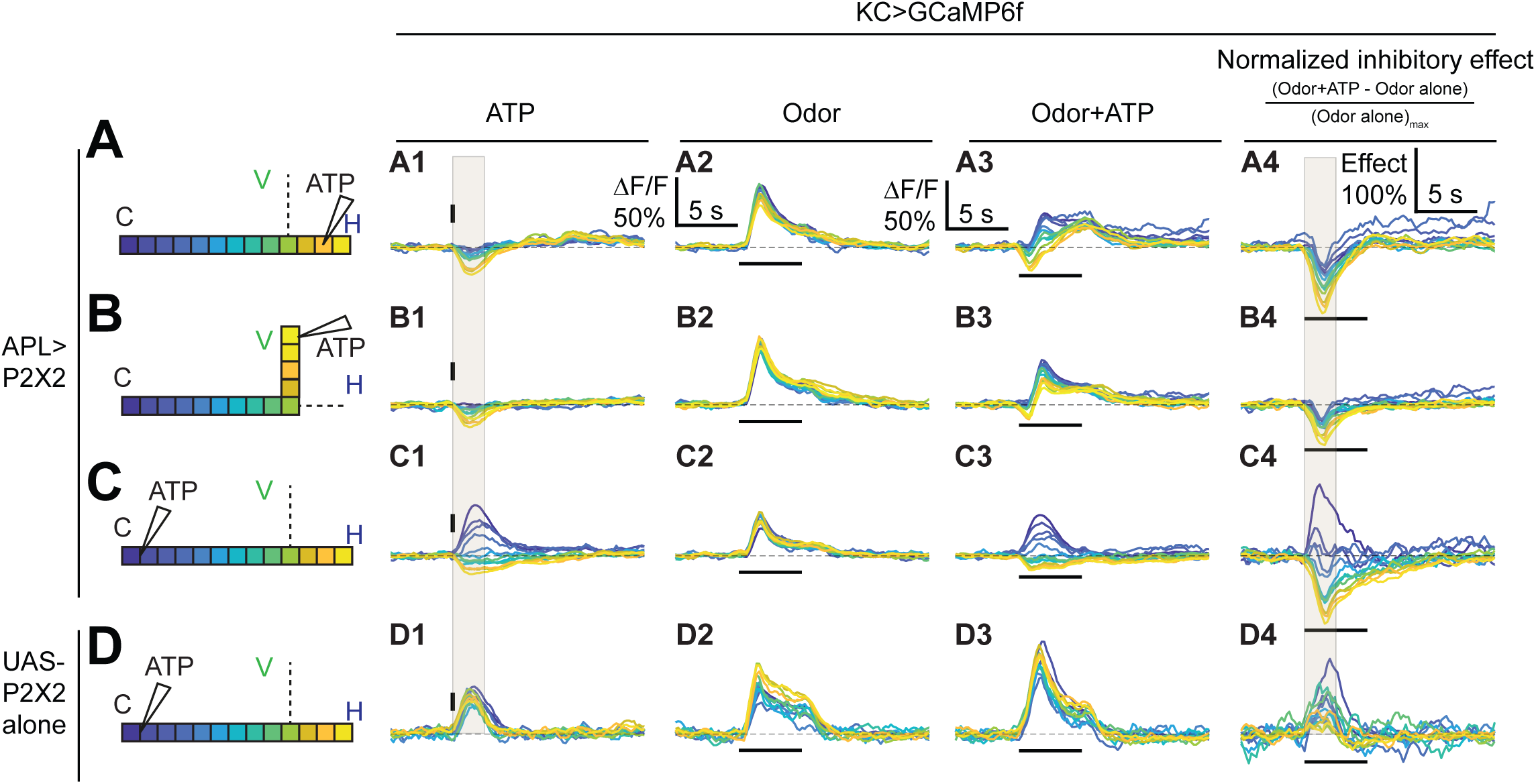
Time courses of APL’s inhibitory effect on KC activity. Traces show the time course of the response in each segment of KCs, averaged across flies, when stimulating APL with 0.75 mM ATP (10 ms puff at 12.5 psi) in the horizontal lobe **(A)**, vertical lobe **(B)**, or calyx **(C). (D)** shows responses in negative control flies (UAS-P2×2 alone). Columns 1-3 show responses of KCs to: **(A1-D1)** activation of the APL neuron by ATP, **(A2-D2)** the odor isoamyl acetate, **(A3-D3)** or both. **(A4-D4)** Normalized inhibitory effect of APL neuron activation on KC responses to isoamyl acetate (i.e., (column 3 - column 2)/(maximum of column 2). The color-coded skeleton indicates which segments have traces shown (dotted lines mean the data is omitted for clarity; the omitted data appear in **Fig. 7**). Vertical and horizontal bars indicate the timing of ATP and odor stimulation, respectively. Gray shading in panels **A1-D1** and **A4-D4** indicate the intervals used for quantification in **Fig. 7A2-C2** and **A4-C4**, respectively. The data in **D1-D4** is quantified in **Fig. S2**. n, given as # neurons (# flies): (A1-A4, B1-B4) 10 (9), (C1-C4) 9 (8), (D1-D4) 5 (3).

**Figure 7.**
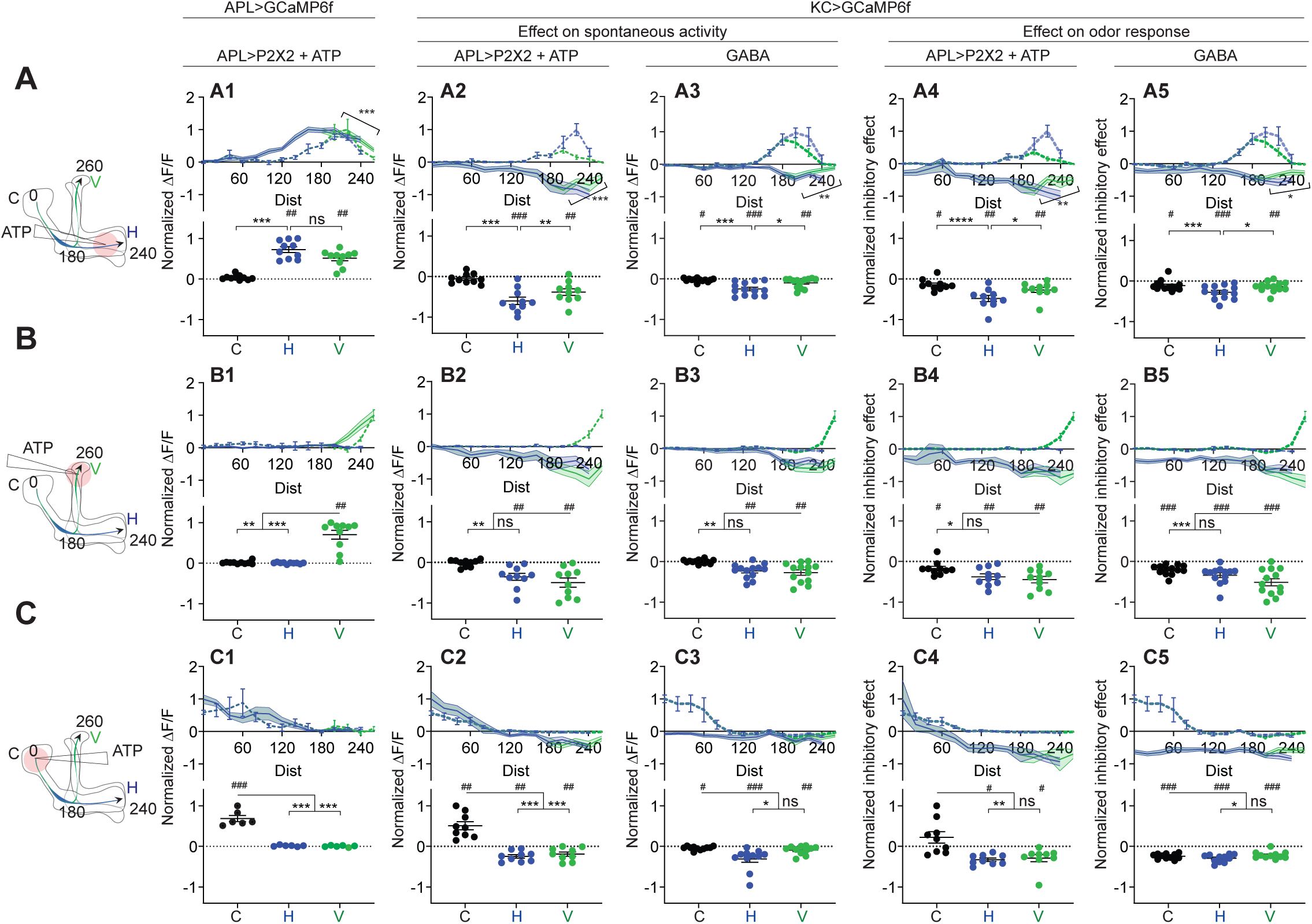
Quantification of the inhibitory effect of GABA or the APL neuron on KC activity. Rows: Local application of ATP (0.75 mM) or GABA (7.5 mM) in the horizontal lobe **(A1-A5)**, vertical lobe **(B1-B5)** or calyx **(C1-C5)**. Columns: Column 1: APL’s response to ATP stimulation **(A1-C1)**, repeated from Fig. 5 for comparison. Columns 2-3: KC responses to local activation of APL by ATP **(A2-C2)** or to GABA application **(A3-C3)**. Columns 4-5: Nnormalized inhibitory effect of APL activation **(A4-C4)** or GABA application **(A5-C5)** on KC responses to isoamyl acetate. Data shown are mean responses in each segment (averaged over time in the gray shaded periods in Fig. 6). The color of the curves matches the vertical (green) and horizontal (blue) lobular branches depicted in the schematics shown on the left. The responses were normalized to the segment (upper panels) or data point (lower panels) with the largest absolute value across matching conditions (columns 2+3, or columns 4+5). Genotypes: for ATP, mb247-LexA>GCaMP6f, APL>P2×2; for GABA, OK107-GAL4>GCaMP6f. The baseline fluorescence for the red dye comes from bleedthrough from the green channel; only trials without odour were used for red dye quantification, in these trials, the change in green bleedthrough (∼10-40%) is negligible compared to the increase in red signal (150-300%). n, given as # neurons (# flies): (A1, B1) 10 (6), (C1) 6 (4), (A2, A4, B2, B4) 10 (9), (C2, C4) 9 (8) (A3, A5, B3, B5) 13 (8), (C3, C5) 11 (6). # p<0.05 ### p<0.001, one-sample Wilcoxon test, or one-sample t-test, vs null hypothesis (0) with Holm-Bonferroni correction for multiple comparisons,. * p<0.05, *** p<0.001, Friedman test with Dunn’s multiple comparisons test, or repeated-measures one-way ANOVA with Holm-Sidak multiple comparisons test, comparing the stimulated site vs the unstimulated sites. Diagonal brackets in (A1-A5), paired t-test or Wilcoxon test comparing the response at segment 200 vs 260 on the vertical branch. See Table S2 for detailed statistics.

**Figure 8.**
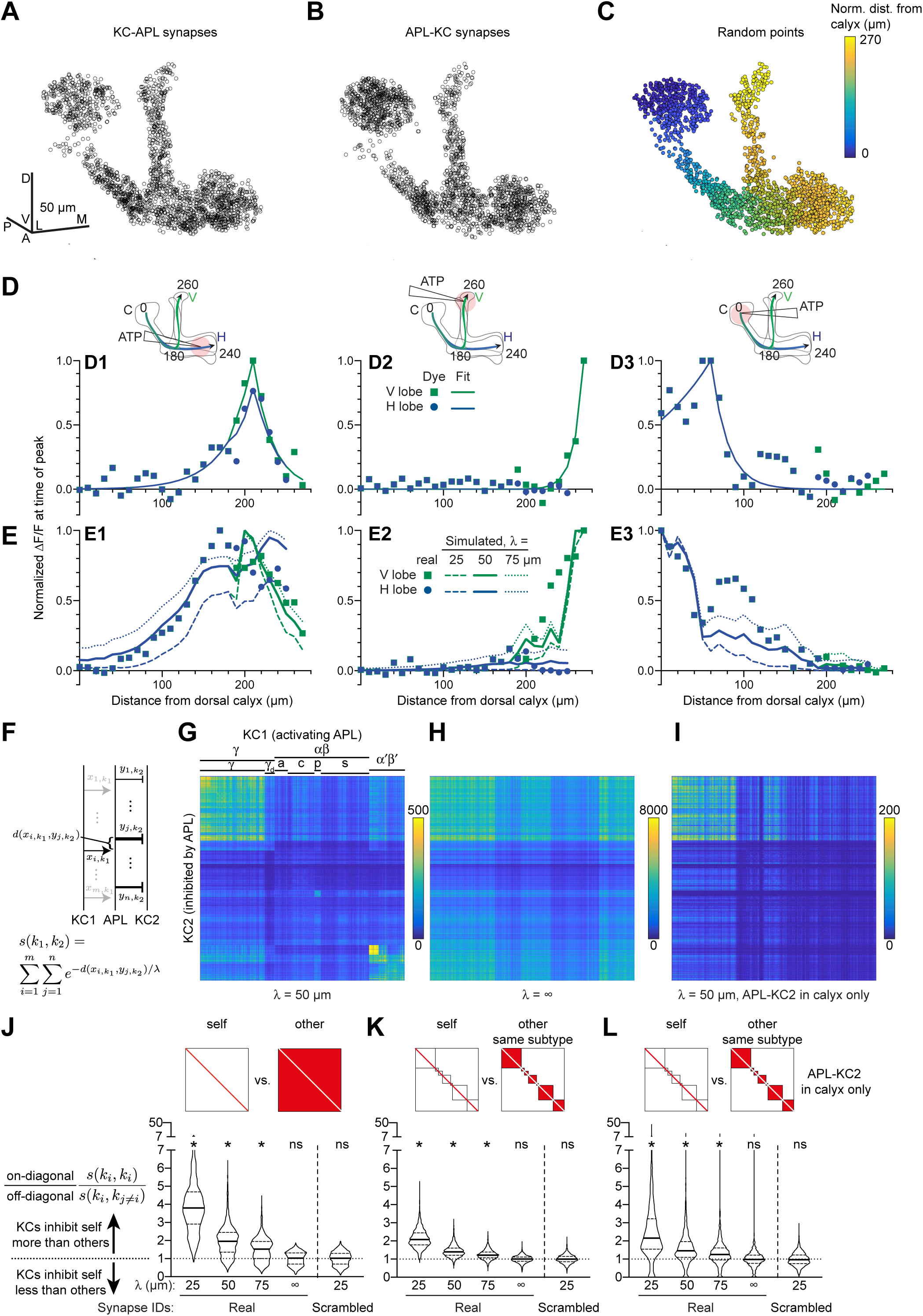
Localized activity and KC-APL anatomy suggest that KCs inhibit themselves more than other KCs. **(A,B)** 2000 randomly selected synapses from the set of 96,866 unique KC-APL synapse locations **(A)** and 15,655 unique APL-KC synapse locations **(B)**. Scale bars 50 µm. The KC-APL synapses in the calyx are consistent with reports of presynaptic release from Kenyon cells in the calyx (Christiansen et al., 2011). The relative lack of KC-APL and APL-KC synapses in the posterior peduncle is consistent with the lack of syt-GFP signal in both APL and Kenyon cells in this zone, thought to be the Kenyon cells’ axon initial segment (Trunova et al., 2011; Wu et al., 2013). **(C)** 2000 random points along APL’s neurite skeleton, color-coded by normalized distance from the dorsal calyx along the average skeleton in **Fig. 5**. **(D)** Red dye signal from **Fig. 5** in 10 µm segments (dots), with fitted exponential decay curves (lines), for stimulation in the horizontal lobe **(D1)**, vertical lobe **(D2)**, and calyx **(D3)**. The slight discontinuity in the curve fit for **D1** arises because the calyx-to-junction branch has slightly different fits for the calyx-to-vertical vs. calyx-to- horizontal branches; where the two merge, we averaged the fits together. **(E)** Real activity in APL from **Fig. 5** (dots) compared to simulated activity for different space constants (lines), given localized stimulation defined by fitted curves in **(D)** in the horizontal lobe **(E1)**, vertical lobe **(E2)**, and calyx **(E3)**. **(F)** Schematic of metric s(k1,k2) for predicting how strongly KC1 inhibits KC2 via APL given the space constant λ and the relative distances (d) between KC1-APL (x_i,k1_) and APL-KC2 (y_j,k2_) synapses along APL’s neurite skeleton. The variable thickness of the APL-KC2 synapses depicts how synapses closer to the KC1-APL synapse (x_i,k1_) are weighted more strongly. The other KC1-APL synapses are greyed out to show that although they are part of the double summation, the focus in the diagram is on distances between x_i,k1_ and APL-KC2 synapses. **(G)** s(k1,k2) for all pairs of KCs, for space constant 50 µm. αβ, α′β′, γ subtypes as annotated in the hemibrain v1.0.1. Note that s(k1,k2) is higher along the diagonal. KCs were sorted by subtype as annotated in the hemibrain dataset, and reordered by hierarchical clustering to place similar KCs within the same subtype next to each other. **(H)** As **(G)** except space constant is infinite. **(I)** As **(G)** except only including APL-KC synapses (y_j,k2_ in panel **F**) that are in the calyx. In **(H-I)**, the order of KCs is preserved from **(G)**. **(J-L)** The violin plots show, for each KC1, the ratio of s(k1,k1) (self-inhibition; on- diagonal pixels in **G-I**) vs. the average of s(k1,k2) (inhibiting other KCs; off-diagonal pixels in **G-I**), given different space constants (λ), where k2 includes all other KCs **(J)**, or all other KCs within the same subtype **(K). (L)** As **(K)** except only including APL-KC synapses in the calyx. The squares above the violin plots mark in red which pixels are being analyzed from a heat map of s(k1,k2) as in panel **G**: i.e., for “self”, the diagonal pixels; for “other”, the off-diagonal pixels; for “other, same subtype”, the off-diagonal pixels for KCs of the same subtype. “Synapse IDs”: For “Scrambled” (but not “Real”), the identities of which Kenyon cell each KC-APL or APL-KC synapse belonged to were shuffled. Solid horizontal lines show the median; dotted lines show 25% and 75% percentile. n = 1921 KCs (8 of 1929 KCs are annotated as atypical or questionable KCs). * p < 0.0001, different from 1.0 by Wilcoxon test with Holm-Bonferroni correction for multiple comparisons.

To test this, we again locally stimulated APL with ATP and used volumetric imaging to record activity in Kenyon cells (expressing GCaMP6f under the control of mb247-LexA) instead of from APL. Although Kenyon cells are mostly silent in the absence of odor, they do exhibit some spontaneous activity (Turner et al., 2008). As expected, stimulating APL with ATP in the lobes reduced the baseline GCaMP6f signal in Kenyon cells, and the inhibitory effect was strongest at the site of APL stimulation. This can be seen in both color-coded time courses (**Fig. 6A1,B1**) and plots of APL’s inhibitory effect vs. distance along the 3D skeleton (**Fig. 7A2,B2**). We saw qualitatively similar results when we examined Kenyon cells’ odor-evoked activity instead of spontaneous activity. Here, we measured the inhibitory effect as the difference between Kenyon cell odor responses with and without local stimulation of APL by ATP, normalized to the response without APL stimulation (**Fig. 6A4,B4**, calculated from panels **A2,B2** and **A3,B3**). Again, this normalized inhibitory effect was strongest at the site of APL stimulation (**Fig. 7A4,B4**). These results provide the first physiological evidence that local activity in APL can locally inhibit Kenyon cells.

Strikingly, local inhibition was frequently stronger proximally in Kenyon cell axons than distally. Specifically, when stimulating APL in the horizontal lobe, odor-evoked Ca^2+^ influx in Kenyon cells was strongly inhibited at the junction of the vertical and horizontal lobes, but less so in the tip of the vertical lobe (**Fig. 7A2, A4**; difference between node 200 µm vs. 260 µm, p < 0.01, paired t-test; see **Table S2**). An analogous difference between 200 vs. 260 µm also occurs in APL activity with the same stimulation (**Fig. 7A1**). That inhibition of Kenyon cell axons can differ over such a short distance suggests that APL can differentially inhibit mushroom body compartments (see Discussion).

Surprisingly, puffing ATP on the calyx excited Kenyon cells in the calyx but inhibited them in the lobes (**Fig. 6C, 7C**). This apparently puzzling result can be explained by observing that in negative control flies with UAS-P2×2 and no GAL4, calyx ATP also excited Kenyon cells, but did not inhibit them (**Fig. 6D, S2**). These results are best explained by leaky expression of P2×2 from the UAS-P2×2 transgene, either in Kenyon cells or projection neurons. The leaky P2×2 expression allows ATP to excite Kenyon cells in their dendrites, driving them to spike (hence ATP drives Ca^2+^ influx throughout Kenyon cell dendrites and axons). However, in APL>P2×2 flies, inhibition from APL is strong enough to suppress Kenyon cell spikes despite the local excitation from leaky P2×2, thereby blocking both spontaneous and odor-evoked activity in Kenyon cell axons. Note that some of the local Ca^2+^ influx in the calyx may reflect Ca^2+^ entering directly through P2×2 channels, despite inhibition from APL shunting away the resulting depolarization. These results show that sufficiently strong local inhibition in Kenyon cell dendrites can silence Kenyon cells.

The inhibitory effect was sometimes less localized than APL activity itself. For example, when ATP was ejected at the tip of the vertical lobe, APL activity decreased to zero by 100 µm from the ejection site and there was no activity in the horizontal lobe (**Fig. 4D,G, 7B1**), whereas the inhibitory effect on Kenyon cells persisted into the peduncle and the horizontal lobe (**Fig. 7B2**), even into the calyx (>200 µm away) in the case of the inhibitory effect on Kenyon cell odor-evoked responses (**Fig. 7B4**). However, this more extended inhibitory effect on the calyx was significantly smaller than the local inhibitory effect on the vertical lobe. These results suggest that APL can, to a modest extent, inhibit areas of Kenyon cells beyond areas where APL itself is active.

### Local GABA application mimics the inhibitory effect of local APL stimulation

How could APL inhibit Kenyon cells in areas where it itself is not active? One explanation could be that because we are measuring GCaMP6f signals and not voltage, APL might be active and thus release GABA even where we do not see increased GCaMP6f signal. This possibility seems unlikely as it is unclear how GABA would be released in the absence of Ca^2+^ influx. Still, to test this possibility, we replaced APL stimulation with direct application of GABA. We reasoned that if local ATP stimulation causes APL to release GABA over a wider region than APL’s own GCaMP6f signal, then directly applying GABA should cause a more spatially restricted inhibitory effect than applying ATP (which would evoke wider release of GABA from APL).

In fact, directly applying GABA caused the same inhibitory effect on Kenyon cells as locally stimulating APL with ATP: strongest at the site of stimulation but still affecting Kenyon cells far from the site of stimulation, to a lesser extent (**Fig. 7**, columns 3 and 5; traces in **Fig. S3**). In particular, GABA applied to the tip of the vertical lobe inhibited Kenyon cells throughout the horizontal lobes. In the case of odor-evoked activity, this inhibition extended to the peduncle and calyx, though significantly weaker than in the vertical lobe (**Fig. 7B3, B5**, compare to **B2, B4**). These results argue against the possibility that APL releases GABA more widely than its area of activity and suggest that APL’s inhibitory effect extends beyond the zone of its own activity, though weaker than where it is active (see Discussion).

In other respects as well, puffing GABA elicited the same effects above noted for locally activating APL with ATP. Locally inhibiting Kenyon cells in the calyx inhibited them throughout the lobes (**Fig. 7C3, C5**, compare to **C2, C4**, now without the confounding effects of leaky P2×2 expression). With local inhibition in the horizontal lobe, Kenyon cells were more inhibited at the junction than at the vertical lobe tip **(Fig. 7A3,A5**, compare to **A2, A4**). The close correspondence of effects of GABA and ATP application indicates that the observed localization of ATP-induced Ca^2+^ influx in APL accurately reflects the localization of GABA release.

### Connectomic analysis predicts that each Kenyon cell disproportionately inhibits itself

We next asked how local inhibition by APL affects lateral inhibition between Kenyon cells. APL is thought to act through all-to-all feedback inhibition to sparsen and decorrelate population odor responses in Kenyon cells (Lin et al., 2014). Such a function assumes that each Kenyon inhibits all Kenyon cells via APL roughly equally and indeed, decorrelation should theoretically work better if all Kenyon cells inhibit each other than if each Kenyon cell inhibits only itself (see Appendix; **Fig. S5**). However, if a Kenyon cell activates APL locally and APL is not evenly activated throughout, it may inhibit other Kenyon cells unevenly, i.e., some more than others. Such uneven lateral inhibition would depend both on how activity spreads within APL and on how APL-KC and KC-APL synapses are spatially arranged on APL neurites. In particular, if activity in APL decays with distance, activating Kenyon cell 1 (KC1) would inhibit another Kenyon cell (KC2) more strongly if KC1-APL synapses are close to APL-KC2 synapses (as measured along APL’s neurites) than if they are far apart. Thus, the degree to which KC1 inhibits KC2 can be described by the weighted count of every pair of KC1-APL and APL-KC2 synapses (see Methods). The weight of each pair should fall off exponentially with the distance between the two synapses, given that cable theory predicts that signals decay as exp(-x/space constant) (Hodgkin and Rushton, 1946) (**Fig. 8F**).

To test whether KC-APL synapses are spatially arranged to allow uneven lateral inhibition, we analyzed the hemibrain connectome dataset released by Janelia FlyEM and Google (v. 1.0.1) (Xu et al., 2020), which contains a fully segmented APL and annotated KC-APL and APL-KC synapses. All 1929 traced Kenyon cells in this volume form reciprocal synapses with APL (49.6±17.9 APL-KC and 52.6±13.4 KC-APL synapses per KC; mean±s.d.). APL-KC synapses (but not KC-APL synapses) are polyadic (a single presynaptic density on APL contacts multiple KCs). We mapped all annotated APL-KC and KC-APL synapses onto APL’s neurite skeleton (181,050 connected nodes, total length 79.5 mm; 95,678 and 101,430 unique synapses mapped to 15,655 and 96,866 unique locations on APL’s skeleton for APL-KC and KC-APL synapses, respectively) (**Fig. 8A,B**) and measured the distance from every APL-KC to every KC-APL synapse along APL’s neurite skeleton.

To determine what space constant to use in our analysis, we mapped our recordings onto the reconstructed APL by dividing it into segments (Voronoi cells) along the mushroom body 3D skeleton as in **Fig. 5-7**. Here we used segments spaced 10 µm apart to increase spatial resolution (in **Fig. 5-7** we used 20 µm spacing to reduce variability across individual flies). We took the red dye signal in each segment from **Fig. 5**, averaged across flies, at the time of the peak signal in the segment receiving the most red dye. Because these signals were weak and noisy (peak ΔF/F ∼20-30%, compared to 300% for GCaMP6f), we fitted exponential decay curves to approximate the ATP stimulus that each segment received. We then simulated **Fig. 5** by placing ∼10,000 randomly distributed points on the APL neurite skeleton (**Fig. 8C**) and “stimulating” each segment of APL according to our curve fits of red dye signal (**Fig. 8D**). Given this localized stimulus, we calculated how much signal an average point in each segment of APL would receive from all other points in APL, given a normalized space constant of 25, 50 or 75 µm (**Fig. 8E**). “Normalized space constant” means the space constant along a neurite of average diameter, i.e. ∼0.45 µm (**Fig. S4D**); we modeled space constants as varying with the square root of the neurite radius (see Methods). Note that for simplicity, we did not consider branching effects in APL. Overall, 50 µm gave the best fit: 25 µm did not spread activity enough, while with 75 µm, activity spread to areas where **Fig. 6** found none **(Fig. 8E)**. Similar results were obtained when modelling activity as decaying along a straight-line mushroom body skeleton that ignores APL’s neurite morphology, albeit with some differences, likely reflecting areas where APL’s neurites do not simply travel parallel to the straight-line skeleton **(Fig. S4C)**.

Using these space constants, our model predicts that Kenyon cells disproportionately inhibit some Kenyon cells more than others (**Fig. 8G-L**). In particular, each Kenyon cell inhibits itself more strongly than it inhibits other individual Kenyon cells on average (**Fig. 8J-L**). (This can be seen in the brighter pixels along the diagonal in **Fig. 8G,I**.) This is due partly to the fact that γ, αβ_p_ and α′β′ Kenyon cells inhibit other Kenyon cells of the same subtype more than Kenyon cells of other subtypes. (This can be seen as the large blocks along the diagonal in **Fig. 8G**). However, even when comparing self-inhibition to inhibition of only other Kenyon cells of the same subtype, each Kenyon cell inhibits itself more than it inhibits other individual Kenyon cells on average (**Fig. 8K**).

These results arise from treating every KC-APL and APL-KC synapse equally. In reality, Kenyon cell spiking is driven by dendritic input in the calyx, as confirmed by our finding that activity in Kenyon cells can be suppressed throughout the mushroom body by locally activating APL or applying GABA in the calyx **(Fig. 6,7**). Therefore, we repeated this analysis, taking into account only APL-KC synapses in the calyx. Again, our model predicts that each Kenyon cell inhibits itself more than it inhibits other individual Kenyon cells on average (**Fig. 8I,L**). Importantly, this imbalance decreases with longer space constants and disappears entirely if the space constant is infinite (at λ = 50 µm, the median imbalance is ∼40%). Moreover, the imbalance arises from the particular spatial relations between synapses of different Kenyon cells, because the imbalance disappeared when we shuffled the identities of which Kenyon cell each KC-APL or APL-KC synapse belonged to **(Fig. 8J-L**). Thus, our physiological measurements of localized activity of APL, combined with the spatial arrangements of KC-APL and APL-KC synapses, predict that Kenyon cells disproportionately inhibit themselves compared to other individual Kenyon cells.

## Discussion

We have shown that activity in APL is spatially restricted intracellularly for both sensory-evoked and local artificial stimulation, consistent with relatively low expression of voltage-gated Na^+^ and Ca^2+^ channels in APL. This local activity in APL translates into localized inhibition onto Kenyon cells. Finally, combining physiological and anatomical data — APL’s estimated space constant and the spatial arrangement of KC-APL and APL-KC synapses – predicts that each Kenyon cell disproportionately inhibits itself more than other individual Kenyon cells.

The locust equivalent of APL is called GGN (‘giant GABAergic neuron’) (Papadopoulou et al., 2011). A compartmental model of GGN predicts that local current injection in the α lobe should lead to depolarization throughout GGN (at least 500 µm across) though the depolarization should decrease with distance from the injection site (Ray et al., 2019). In contrast, we found that local stimulation of APL with P2×2+ATP decayed to undetectable levels within as little as 100 µm. What may explain this difference? Although GGN and APL are both widefield inhibitory interneurons in the mushroom body, they have striking morphological differences. For example, GGN separately innervates the lobes and calyx with numerous thin, branching processes; these tangled innervations are joined by relatively thick processes (almost 20 µm in diameter) (Ray et al., 2019). In contrast, APL innervates the lobes continuously with numerous neurites that emerge in parallel from the calyx (Mayseless et al., 2018; Xu et al., 2020). In the FIB-SEM hemibrain (Xu et al., 2020), these neurites average ∼0.4-0.5 µm in diameter throughout APL; ∼95% of neurite length is less than 1 µm in diameter and the maximum diameter is ∼3 µm (**Fig. S4D**). GGN’s morphology actually more resembles *Drosophila*’s larval APL, which has separate innervations of the lobes and calyx joined by a single process (Eichler et al., 2017; Masuda-Nakagawa et al., 2014). The larval APL has pre-synapses only in the calyx, while adult APL has pre-synapses everywhere (**Fig. 8A**). It is tempting to speculate that the locust and larval *Drosophila* mushroom bodies use global activity in APL/GGN to send feedback from Kenyon cell axons to Kenyon cell dendrites, whereas the adult *Drosophila* mushroom body uses local activity in APL to locally inhibit Kenyon cells everywhere. The larval APL might be small enough for activity to spread passively from lobes to calyx (∼100 µm) or might have different ion channel composition from adult APL; the large diameter of the locust GGN’s neurites between lobe and calyx might increase their space constants enough to allow activity to propagate.

We measured APL’s local effects by locally puffing ATP or GABA on the mushroom body. How spatially precise was this application? The co-ejected red dye indicator spread tens of microns from the ejection site (decay to half-strength ∼10-25 µm; see Table S2). This spatial resolution is close to that of other techniques for local *in vivo* stimulation like two-photon optogenetic stimulation (decay to half-strength 12 µm horizontal, 24 µm axially: (Packer et al., 2015)). How precisely does the red dye indicate the spread of the bioactive compound (ATP/GABA)? This is difficult to answer with certainty, as it is unclear how much of the spread of dye/ATP/GABA arises from bulk flow from ejection from the pipette (which should affect dye, ATP and GABA equally) vs. diffusion (which depends on molecular weight and volume). To the extent that ATP or GABA spread differently from the red dye, they are likely to spread farther, as they have lower molecular weights (dye: 1461; ATP: 507; GABA: 103), meaning that our stimulation is less spatially restricted than the dye suggests. Thus, our results likely overestimate, rather than underestimate, how far APL activity and APL’s inhibitory effect spread from the site of stimulation. If the space constant is indeed lower than our normalized estimate of 50 µm, then the imbalance between Kenyon cells inhibiting themselves vs. others is likely even more severe (compare λ = 25 µm and 50 µm on **Fig. 8J-L**). In any case, in comparing the spatial spread of activity in APL vs. APL’s inhibitory effect, we used the same protocol for ATP stimulation, so these results are directly comparable even if the exact distance of ATP spread is uncertain.

We showed that APL can locally inhibit Kenyon cells in multiple locations in the mushroom body. Remarkably, inhibition of Ca^2+^ influx in Kenyon cell axons could be stronger proximally than distally (**Fig. 7A**), as though action potentials traveled through a zone of inhibition and recovered on the other side. This was true both when stimulating APL with ATP or when directly applying GABA. GABA should act on Kenyon cells by suppressing depolarization through shunting inhibition in both dendrites and axons, as the GABA_A_ receptor Rdl is expressed in both (Liu et al., 2007). Yet if action potentials are blocked proximally, how can they recover distally? It may be that although depolarization is suppressed in a zone of local inhibition, enough current remains on the other side to regenerate the action potential. Alternatively, local inhibition might particularly suppress Ca^2+^ influx rather than depolarization, perhaps by acting through GABA_B_ receptors; Kenyon cells express GABA_B_R1 and GABA_B_R2 (Aso et al., 2019; Crocker et al., 2016; Croset et al., 2018; Davie et al., 2018), although their subcellular localization is unknown. Given that synaptic vesicle release requires Ca^2+^ influx, such inhibition would still suppress Kenyon cell synaptic output.

While APL inhibits Kenyon cells locally, the inhibitory effect spreads somewhat more widely than APL’s own activity, though weakly (**Fig. 7A4-5,B4-5**). Local GABA application produces similar results to locally activating APL with ATP (**Fig. 7**), suggesting that GCaMP6f signals in APL accurately predict GABA release. How can APL inhibit Kenyon cells where it itself is not active? Wider inhibition when inhibiting Kenyon cell dendrites (**Fig. 7C**) can be easily explained as blocking action potentials, but the wider inhibition when inhibiting the axons is more puzzling. This might arise from wider network activity. For example, Kenyon cells form recurrent connections with DPM and dopaminergic neurons and form synapses and gap junctions on each other (Liu et al., 2016; Takemura et al., 2017). Through such connections, an odor-activated Kenyon cell might excite a neighboring Kenyon cell’s axon; the neighbor might passively spread activity both forward and backward or, not having fired and thus not being in a refractory period, it might even be excited enough to propagate an action potential back to the calyx. Alternatively, Kenyon cells indirectly excite antennal lobe neurons in a positive feedback loop (Hu et al., 2010). In these scenarios, locally puffing GABA or activating APL artificially in the vertical lobe tip would block the wider network activity, thus reducing Ca^2+^ influx in the calyx. Although the calyx was not activated when stimulating Kenyon cells in the vertical lobe tip in **Fig. 4B**, this might be because simultaneous activation of all Kenyon cells (but not odor-evoked activation of ∼10% of Kenyon cells) drives strong-enough local APL activation to block wider network activity.

By combining our physiological measurements of localized activity with the detailed anatomy of the FIB-SEM hemibrain connectome (Xu et al., 2020), we built a model that predicts that the average Kenyon cell inhibits itself more than it inhibits other individual Kenyon cells. This prediction is supported by previous experimental results that some Kenyon cells can inhibit some Kenyon cells more than others, and that an individual Kenyon cell can inhibit itself (Inada et al., 2017). Our model goes beyond these results in predicting that the average Kenyon cell actually *preferentially* inhibits itself (**Fig. 8**), and by explaining how differential inhibition can arise from local activity in APL and the spatial arrangement of KC-APL and APL-KC synapses.

We based our phenomenological model on the heuristic that depolarization decays exponentially with distance. The model is constrained by experimental data, though these are necessarily imperfect; potential sources of error include automated segmentation and synapse annotation in the connectome, imperfect registration of the hemibrain APL to our average 3D skeleton, and noisy red dye signals that may not perfectly report ATP localization (see above). We ignored certain complicating factors in our model for simplicity – some due to lack of data (e.g., active conductances; differences in synaptic strength or membrane conductance across APL), others as simply beyond the scope of this study (e.g., dynamic effects of feedback inhibition; effects of neurite branching and differing branch lengths). However, these simplifications and potential error sources would lead us to underestimate, not overestimate, spatial differences in APL activity (and therefore the bias toward self-rather than other-inhibition in Kenyon cells). Future detailed compartmental modeling will allow further insights into local feedback inhibition in the mushroom body.

Our findings show that localized activity within APL has two broader implications for mushroom body function. First, local activity in APL leads to local inhibition of Kenyon cells, to the point that different compartments of the mushroom body lobes can receive different inhibition. This local inhibition would allow the single APL neuron to function effectively as multiple inhibitory interneurons, each locally modulating the function of one mushroom body ‘compartment’, i.e., a unit of dopaminergic neurons / Kenyon cells / mushroom body output neurons. Different compartments are innervated by different dopaminergic neurons (e.g., reward vs. punishment) and different mushroom body output neurons (e.g., avoid vs. approach) (Aso et al., 2014), and they govern synaptic plasticity and hence learning by different rules (e.g., different speeds of learning/forgetting) (Aso and Rubin, 2016). Thus, APL could locally modulate different compartment-specific aspects of olfactory learning. If such local inhibition is important for learning, the fact that spatial attenuation of APL’s inhibitory effect is gradual and incomplete could explain why mushroom body compartments are arranged in their particular order, with reward and punishment dopaminergic neurons segregated into the horizontal and vertical lobes, respectively. Under this scenario, APL would serve two distinct, spatially segregated functions: enforcing Kenyon cell sparse coding in the calyx, and modulating learning in the compartments of the lobes.

Second, the finding of strong self-inhibition compared to other-inhibition provides a new perspective on APL’s function. Inhibition from APL sparsens and decorrelates Kenyon cell odor responses to enhance learned odor discrimination (Lin et al., 2014) but in general, decorrelation is better served by all-to-all lateral inhibition than by self-inhibition (see Appendix, **Fig. S5**), raising the question of why self-inhibition is so strong. Self-inhibition does help decorrelate population activity by pushing some neurons’ activity below spiking threshold (**Fig. S5**); this could occur if APL can be activated by subthreshold activity in Kenyon cells, e.g., from KC-APL synapses in Kenyon cell dendrites. However, this effect of self-inhibition is better thought of, not as decorrelation per se, but rather as gain control (Asahina et al., 2009; Olsen et al., 2010; Root et al., 2008), effectively equivalent to adjusting the threshold or gain of excitation according to the strength of stimulus. Beyond its role in sparse coding, APL inhibition is also thought to function as a gating mechanism to suppress olfactory learning (Liu and Davis, 2008; Zhou et al., 2019); for such a function it would make sense for Kenyon cells to disproportionately inhibit themselves. Future work will address how lateral inhibition interacts with other functions for APL in this local feedback circuit.

## Methods

### Fly strains and husbandry

Flies were kept at 25 °C (experimental) or 18 °C (long-term stock maintenance) on a 12 h/12 h light/dark cycle in vials containing 80 g medium cornmeal, 18 g dried yeast, 10 g soya flour, 80 g malt extract, 40 g molasses, 8 g agar, 25 ml 10% nipagin in ethanol, and 4 ml propionic acid per 1 L water.

VT43924-Gal4.2 was created via LR Clonase reaction between pENTR-VT43924 and pBPGAL4.2Uw-2, in which the Gal4 sequence has been codon-optimized for *Drosophila* and the *hsp70* terminator was replaced with the SV40 terminator (Pfeiffer et al., 2010). The resulting construct was inserted in *attP2* by the University of Cambridge Fly Facility Microinjection Service. lexAop-P2×2 was created by subcloning P2×2 from pUAST-P2×2 (Lima and Miesenböck, 2005) into pLOT (Lai and Lee, 2006). QUAS-GCaMP3 was created by subcloning GCaMP3 (Tian et al., 2009) into pQUAST (Potter et al., 2010). pLOT-P2×2 and pQUAS-GCaMP3 were injected by Genetic Services, Inc.

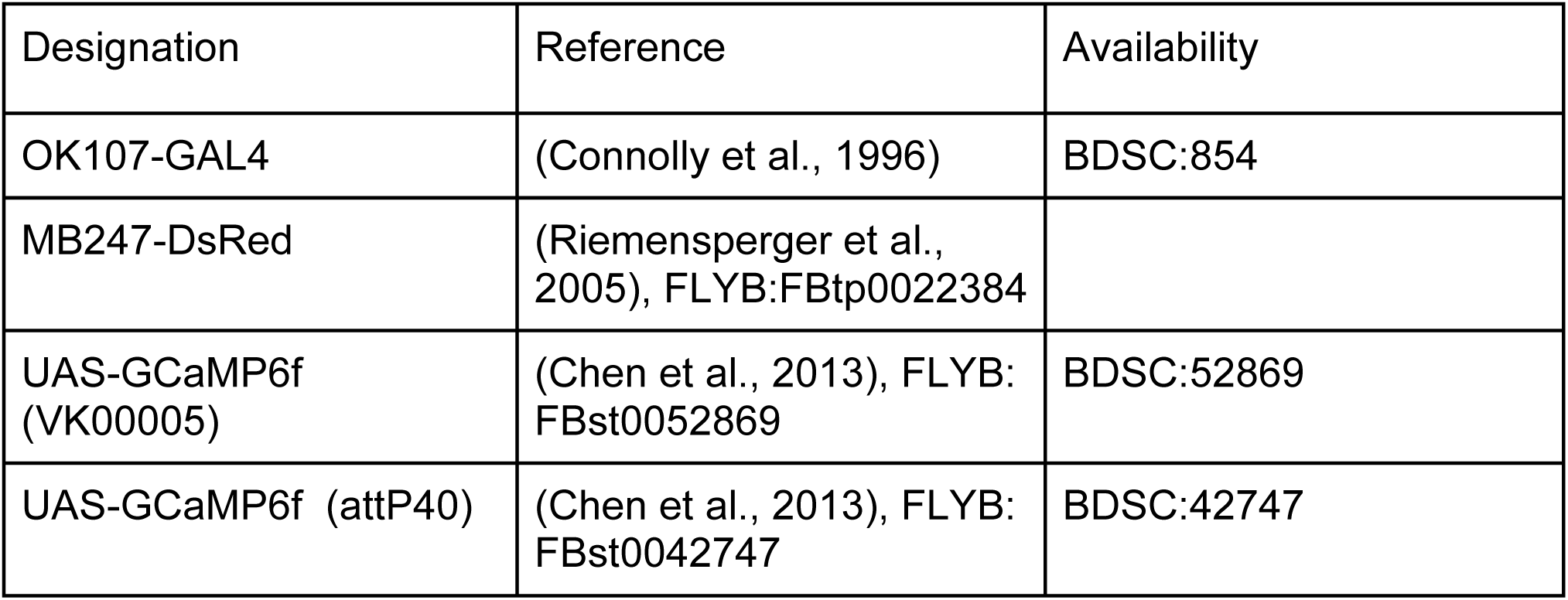
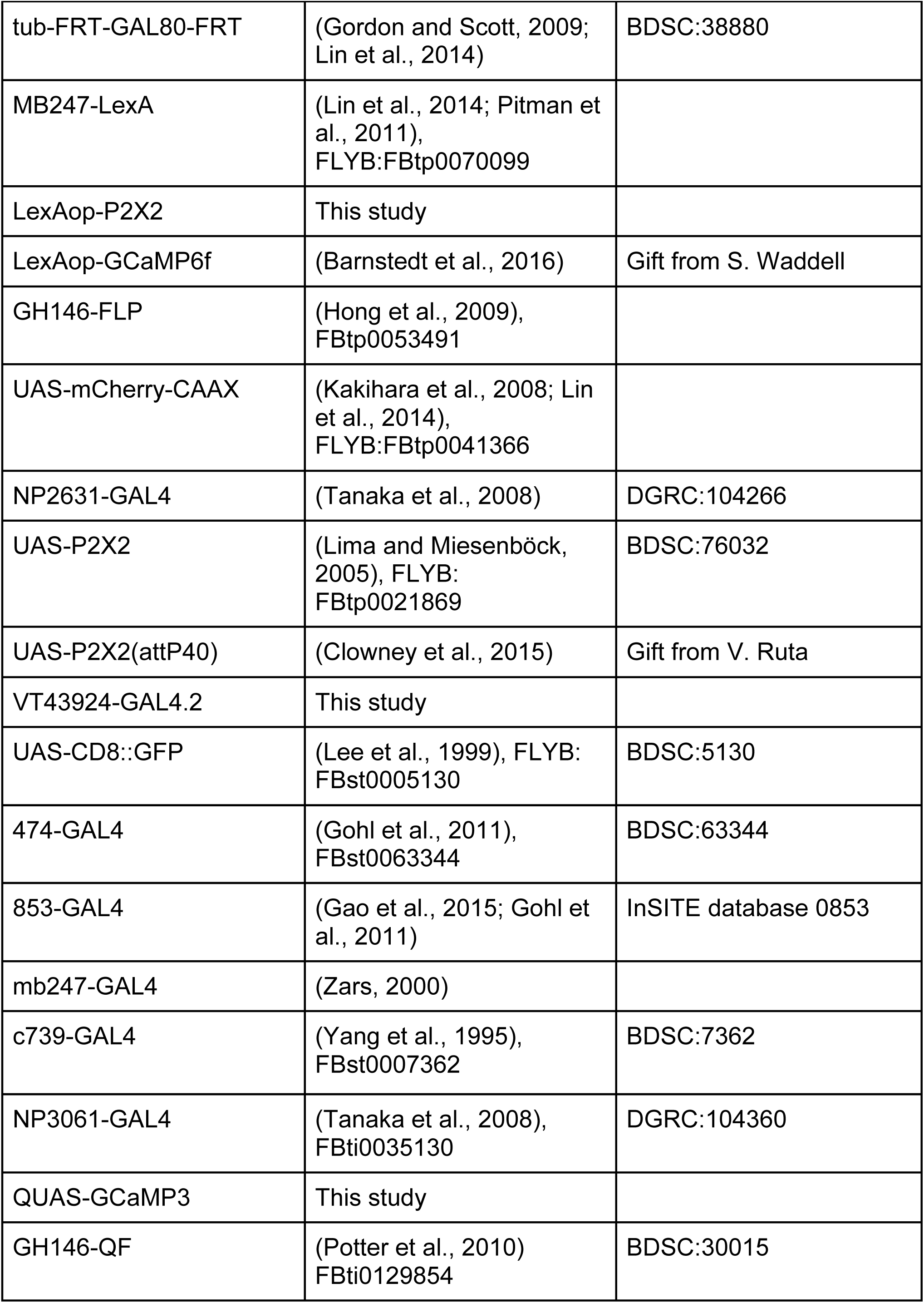

### Odor delivery

Odors were delivered by switching mass-flow controlled carrier and stimulus streams (Sensirion) via solenoid valves (The Lee Company) controlled by LabVIEW 2015 software (National Instruments). Air flow was 0.5 l/min at the exit of the odor tube, which was positioned ∼1 cm from the fly’s head. Odor pulses lasted 5 s. At the end of each recording where odor pulses were given, the tubes were cleaned by passing pure air through. A tube connected to a vacuum pump was positioned behind the fly to deplete the air of odors remaining after the odor pulse.

### Drug delivery

ATP or GABA at the concentrations indicated in the figure legends, together with a red dye to visualize fluid ejection (SeTau-647, SETA BioMedicals), were applied locally by pressure ejection from patch pipettes (resistance ∼10 MΩ; capillary inner diameter 0.86 mm, outer diameter 1.5 mm - Harvard Apparatus 30-0057) coupled to a Picospritzer III (Parker) (puff duration 10 ms, pressure 12.5 psi unless otherwise indicated), which was triggered by the same software as odor delivery. The patch pipette was mounted on a micromanipulator (Patchstar, Scientifica) and positioned at the tip of the vertical lobe, close to the junction point in the horizontal lobe, or in the central part of the calyx. Before initiating recording, fluid was manually ejected to ensure that the pipette was not blocked. For experiments where ATP or GABA were applied during odor stimulus, the fluid ejection was triggered at the same time as the odor pulse, although the odor arrived slightly later due to the travel time of the air stream.

### Electric shock

A rectangle of stacked copper plates was brought into contact with the fly’s abdomen using a manually movable stage (DT12XYZ/M, Thorlabs). The design of the copper plates was based on (Felsenberg et al., 2018). Shocks were provided by a DS3 Constant Current Isolated Stimulator (1.2 s, 32 mA, Digitimer; maximum voltage 90 V) and triggered by the same software as the odor delivery. Successful shock delivery was confirmed by observing the fly’s physical reaction with a Genie Nano-M1280 camera (Teledyne Dalsa) coupled to a 1x lens with working distance 67 mm (SE-16SM1, CCS).

### Functional imaging

Flies were cold-anesthetized and mounted using wax and dental floss in a hole in a piece of aluminum foil fixed to a perfusion chamber, such that the fly’s dorsal and ventral sides were kept on opposite sides of the foil. The dorsal part was immersed in carbogenated (95% O_2_, 5% CO_2_) external solution (103 mM NaCl, 3 mM KCl, 5 mM trehalose, 10 mM glucose, 26 mM NaHCO_3_, 1 mM NaH_2_PO_4_, 3 mM CaCl_2_, 4 mM MgCl2, 5 mM TES, pH 7.3). The cuticle overlying the brain was carefully removed using forceps, followed by removal of fat tissue and trachea. During experiments, the brain was continuously perfused with carbogenated external solution (1.96 ml/min) using a Watson-Marlow pump (120S DM2).

Brains were initially inspected using widefield microscopy (Moveable Objective Microscope, Sutter) and a xenon-arc lamp (LAMBDA LS, Sutter). Functional imaging was carried out using two-photon laser-scanning microscopy (Ng et al., 2002; Wang et al., 2003). Fluorescence was excited by a Ti-Sapphire laser (Mai Tai eHP DS, Spectra-Physics; 80 MHz, 75-80 fs pulses) set to 910 nm, which was attenuated by a Pockels cell (350-80LA, Conoptics) and steered by a galvo-resonant scanner (RESSCAN-MOM, Sutter). Excitation was focussed using a 1.0 NA 20X objective (XLUMPLFLN20XW, Olympus). Emitted light passed through a 750 nm short pass filter (to exclude excitation light) and bandpass filters (green: 525/50; red: 605/70), and was detected by GaAsP photomultiplier tubes (H10770PA-40SEL, Hamamatsu Photonics) whose currents were amplified (TIA-60, Thorlabs) and transferred to the imaging computer, which ran ScanImage 5 (Vidrio). Volume imaging was carried out using a piezo objective stage (nPFocus400, nPoint).

Results in **Fig. 3** were generated similarly but on different equipment as described in (Lin et al., 2014): Movable Objective Microscope with galvo-galvo scanners, Chameleon Ultra II laser (Coherent), 20x, 1.0 NA W-Plan-Apochromat objective (Zeiss), HCA-4M-500K-C current amplifier (Laser Components), MPScope 2.0 software (Sutter). Flies were heated with a TC-10 temperature controller (NPI) and HPT-2 inline perfusion heater (ALA). The temperature at the fly was measured with a TS-200 temperature sensor (NPI) and a USB-1208FS DAQ device (Measurement Computing) at 30 Hz; temperature traces were smoothed over 20 frames by a moving-average filter to remove digitization artifacts.

### Structural imaging

Dissected VT43924-GAL4.2>GFP brains were fixed in 4% (wt/vol) paraformaldehyde in PBT (100 mM Na_2_PO_4_, 0.3% Triton-X-100, pH 7.2), washed in PBT (2 quick washes, then 3 20 min washes) and mounted in Vectashield. Confocal stacks were taken on a Nikon A1 confocal microscope in the Wolfson Light Microscopy Facility.

### Image analysis

Motion correction was carried out as in (Lin et al., 2014); flies with excessive uncorrectable motion were discarded. For experiments without 3D-skeletonization (see below), ΔF/F was calculated in ImageJ for manually drawn regions of interest (ROIs). Background fluorescence was taken from an empty region in the deepest z-slice, and subtracted from the ROI fluorescence. F_0_ was calculated as the average fluorescence during the pre-stimulus period and ΔF/F was calculated as (F-F_0_)/F_0_. ΔF/F data were smoothed by a boxcar filter of 5 frames (**Fig. 3**) or 1 s (**Fig 1, 4, 5C-E, 6, 8**) for displaying time course traces or calculating the peak response, but not for calculating the average response (**Fig. 5F-H, 7**).

### 3D-skeletonization

Where the same neuron was recorded in multiple conditions, movies were aligned by maximizing cross-correlation between the time-averaged mb247-dsRed or mb247-LexA>GCaMP6f signals of each recording. This aligned image was used to define the 3D volume of the mushroom body neuropil by manually outlining it in each z-slice (excluding Kenyon cell somata). This volume mask was converted to a skeleton passing through the center of the major branches of the mushroom body (vertical lobe, horizontal lobe, peduncle/calyx), by manually defining key points (e.g., tip of the vertical/horizontal lobes, junction, places where the structure curves) and connecting them into an undirected graph.

To compare skeletons across different flies, a ‘standard’ APL skeleton was generated by taking the average length of each branch across all flies (**Fig. 5**). Each individual skeleton was normalized to this standard skeleton by placing nodes at spacing x*20 µm, where x is the ratio of the branch length of the individual skeleton over the branch length of the standard skeleton. The 3D volume of each APL neuron was partitioned into Voronoi cells, each with one of the evenly spaced skeleton nodes as the centroid. For each node, fluorescence signal at each time point was averaged over all voxels in the corresponding Voronoi cell. The background was subtracted and baseline fluorescence calculated as the average fluorescence during the pre-stimulus period. ΔF/F was calculated for each node and time point as the difference between the fluorescence at that time point and the baseline fluorescence, divided by the baseline fluorescence.

ΔF/F at each node was calculated as both the time-average over the stimulus period (for individual flies), and as the average across flies within a group, to show the time course. In the latter case, time courses were linearly interpolated to a frame time of 0.2 s. ΔF/F vs. distance was plotted according to the normalized distance of each node from the dorsal calyx on the ‘standard’ skeleton. Individual nodes were excluded if the standard deviation of ΔF/F during the pre-stimulus period was greater than 1.0 (green channel), or 0.7 (red channel), as such high noise indicated poor signal in that segment. Normalized inhibitory effect (of ATP or GABA) was calculated as (ΔF/F odor with ATP/GABA - ΔF/F odor alone) divided by the peak ΔF/F for odor alone. We divided by peak rather than average odor response because this was more robust for recordings where the odor response was low, where normalizing to the average caused extreme magnification of small changes.

### Connectome analysis

We reasoned that the strength of lateral inhibition of one Kenyon cell (KC1) onto another (KC2) could be modelled as

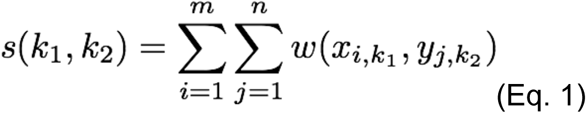

where x_i,k1_ is the ith synapse from KC1 onto APL, and y_j,k2_ is the jth synapse from APL onto KC2, and w is given by

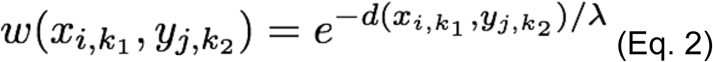

where d is the distance between the two synapses along the APL neurite skeleton and λ is the space constant. To take into account the varying widths of APL neurites, we divided the Euclidean distance along each segment of the APL neurite skeleton by the square root of the radius of that segment (in pixels). This approximates the rule that space constants vary with the square root of the radius, so a wider segment is ‘shorter’ (in electrotonic distance) than a narrower segment. The ‘electrotonic’ total length of the APL skeleton was 16.8 mm compared to the actual length of 79.5 mm, so we divided the space constant in real distance by ∼4.73 to apply it to ‘electrotonic’ distances.

To calculate the distance from every KC-APL synapse to every APL-KC synapse (setting no lower threshold on confidence of synapse annotation), we calculated the distance from each APL-KC synapse to its nearest neighbor APL-KC synapses, as measured along the skeleton of APL neurites. We used Dijkstra’s algorithm on the resulting undirected graph to calculate the distance from each APL-KC synapse to every other APL-KC synapse. We then calculated the distance from each KC-APL synapse to its nearest neighbor APL-KC synapses, and combined this with the distances between APL-KC synapses to find the distance from every KC-APL synapse to every APL-KC synapse.

To simulate the space constant, we placed ∼10,000 points at random on APL’s neurite skeleton. We divided the mushroom body skeleton (**Fig. 6**) into 10 µm segments. We simulated the activity in segment s_i_ (i = 1…35) as the count of every pair between points in s_i_ and points in the whole APL, weighted by the distance separating the pair:

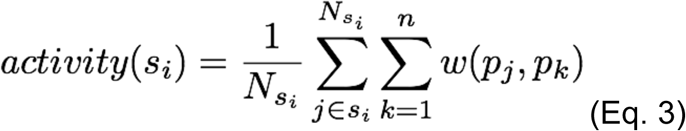

where w(p_j_, p_k_) depends on the distance between points p_j_ and p_k_ (as in Eq. 2), N_si_ is the number of points (j∈s_i_) lying within segment s_i_, and n = the number of random points (∼10,000). We excluded points outside the mushroom body (e.g., the neurite leading to the APL soma).

### Analysis of RNA-seq data

Raw transcripts per million (TPM) counts (Aso et al., 2019) were summed arithmetically across splice variants for each gene. The log_10_ of each resulting value was taken. To prevent negative infinities, we set the log_10_ of zero values to be -2 (i.e., TPM = 0.01), because the minimum TPM value in the dataset was 0.1. The arithmetic mean of log_10_(TPM) was taken across biological replicates for each cell type (i.e., geometric mean of raw TPM). Gene Ontology (Ashburner et al., 2000; The Gene Ontology Consortium, 2019) was used to categorize by biological process the 1529 genes in which APL’s TPM counts were higher than all 20 other cell types in the dataset. The list of ion channels was curated from The Interactive Fly and (Groschner et al., 2018); genes whose mean TPM across all non-APL MB neurons was less than 1 were excluded.

### Statistics and software

Statistical analysis and curve fitting were carried out in Prism 8 (GraphPad). Parametric (t-test, ANOVA) or non-parametric tests (Wilcoxon, Mann-Whitney, Friedman, Kruskal-Wallis) were used depending on whether data passed the D’Agostino-Pearson (or Shapiro-Wilk, for small n) normality test. Traces of manual ROIs were analysed in Igor Pro 7 (WaveMetrics). 3D-skeletonisation and analysis of connectome and RNA-seq data was carried out with custom software written in Matlab (MathWorks). No explicit power analysis was used to pre-determine sample sizes; we used sample sizes comparable to those used in similar studies (e.g., (Zhou et al., 2019)). The experimenter was not blind to experimental conditions or genotypes.

## Acknowledgements

We thank Moshe Parnas, Przemyslaw Stempor, Natalia Bulgakova, and members of the Lin and Juusola labs for discussions, Yoshinori Aso for sharing the RNA-seq data before publication, Saket Navlakha for help with accessing the hemibrain data, Mark Kelly for preliminary analyses of data from (Takemura et al., 2017), Emmanuel Perisse for the design of the electric shocker, Vanessa Ruta, the Bloomington Stock Center, and the Vienna Drosophila Resource Center for fly stocks, and Lily Bolsover, Chloe Donahue, Kath Whitley, Josh Marston, Rachael Thomas, and Rachid Achour for technical assistance. This work was supported by the European Research Council (639489) and the Biotechnology and Biological Sciences Research Council (BB/S016031/1).

## Author contributions

H.A.: Conceptualization, Software, Formal analysis, Investigation, Visualization, Methodology, Writing–original draft, Writing–review and editing

R.S.-G.: Investigation, Writing–review and editing

E.V.: Resources, Writing–review and editing

A.C.L.: Conceptualization, Software, Formal analysis, Supervision, Funding acquisition, Investigation, Visualization, Methodology, Writing–original draft, Writing– review and editing

## Figure legends

**Figure S1.**
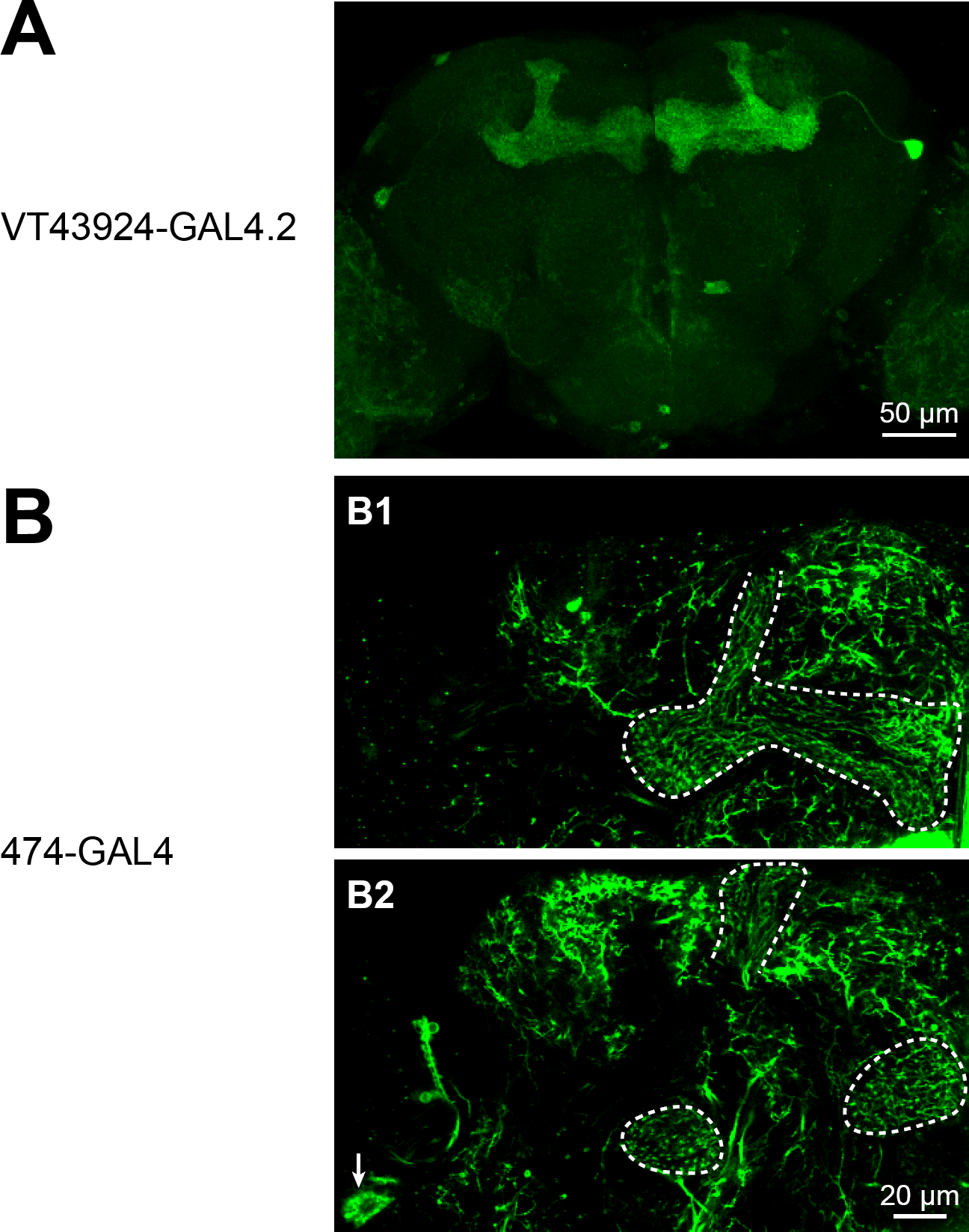
Expression patterns of driver lines. **(A)** Maximum intensity projection of confocal image of VT43924-GAL4.2-SV40 > UAS-CD8::GFP. **(B)** Individual z-slices of 474-GAL4 > UAS-CD8::GFP (unfixed brain imaged on two-photon). B1 shows APL in mushroom body lobes (dashed white outline); B2 shows APL in the anterior peduncle, tip of the vertical lobe, and tip of the β′ lobe (dashed white outlines) and the APL cell body (arrow). B1 and B2 are 18 µm apart. Note the typical morphology of APL with neurites running largely parallel to Kenyon cells. A complete z-projection of 474-GAL4’s expression pattern is available at http://www.columbia.edu/cu/insitedatabase/adultbrain/IT.0474.jpg (Gohl et al., 2011). mb247-GAL4, c739-GAL4, NP3061-GAL4 and GH146-QF were described previously (Potter et al., 2010; Qin et al., 2012; Zhou et al., 2019). The expression pattern of 853-GAL4 is available at http://www.columbia.edu/cu/insitedatabase/adultbrain/IT.0853.jpg

**Figure S2.**
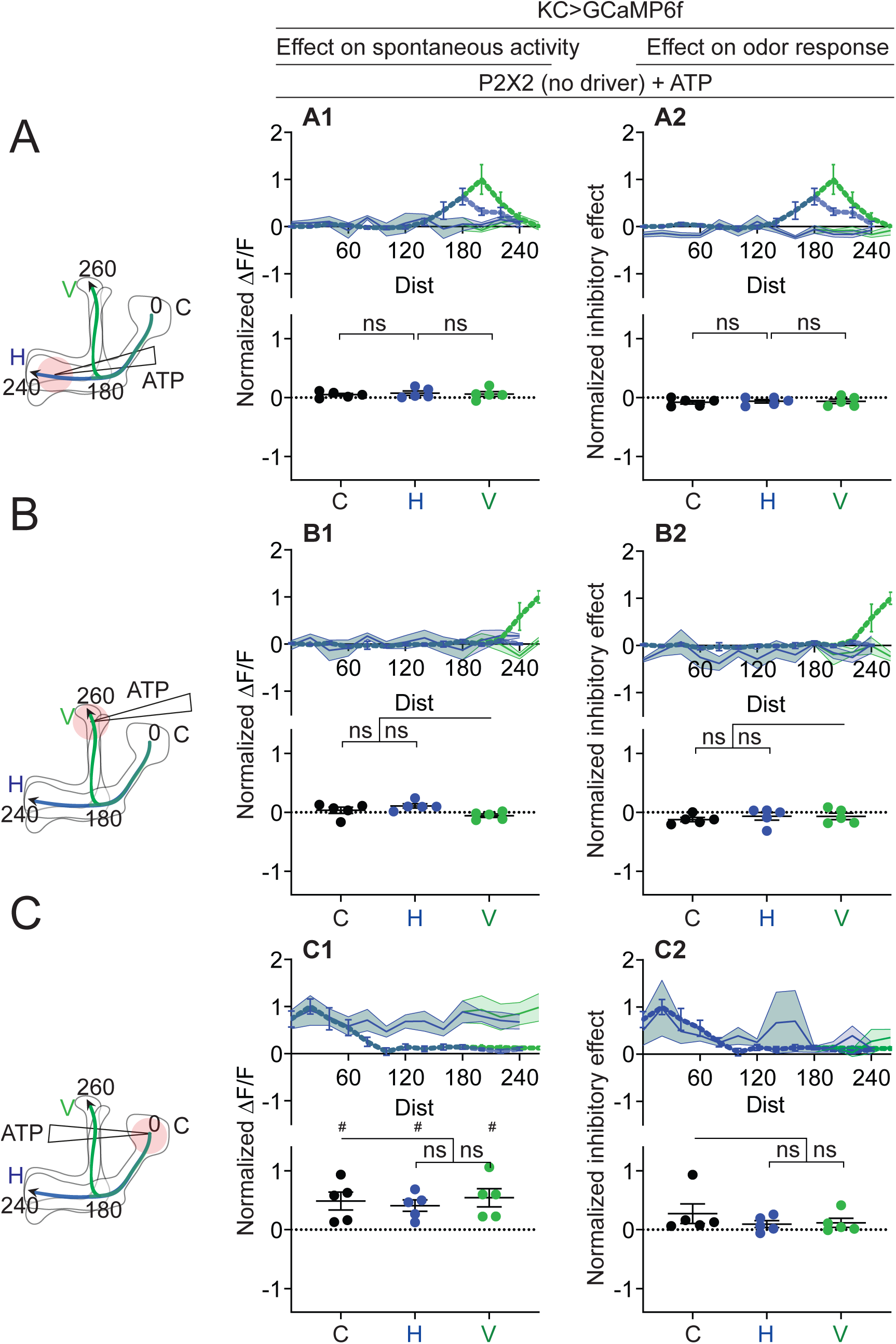
Additional data for Fig. 7: quantification of effect of ATP on negative control (UAS-P2×2 only) Rows: Local application of ATP (0.75 mM) in the horizontal lobe **(A1-A2)**, vertical lobe **(B1-B2)** or calyx **(C1-C2)**. Columns: KC responses to APL activation by ATP **(A1-C1)**, or the normalized inhibitory effect of APL neuron activation on KC responses to isoamyl acetate **(A2-C2)**. Data shown are mean responses in each segment (averaged over time in the gray shaded periods in Fig. 6). The color of the curves matches the vertical (green) and horizontal (blue) lobular branches depicted in the schematics shown on the left. The responses were normalized to the segment (upper panels) or data point (lower panels) with the largest absolute value in the corresponding experimental conditions (columns 2 and 3 in **Fig. 7** for column 1; columns 4 and 5 in **Fig. 7** for column 2). N: 5 neurons, 3 flies. # p<0.05, one-sample t-test, vs null hypothesis (0) with Holm-Bonferroni correction for multiple comparisons. ns, p>0.05, one-way ANOVA with Holm-Sidak’s multiple comparisons test, comparing the stimulated site vs the unstimulated sites. Paired t-test comparing the response at segment 200 vs 260 on the vertical branch gives p=0.2218 (A1), p=0.9397 (A2).

**Figure S3.**
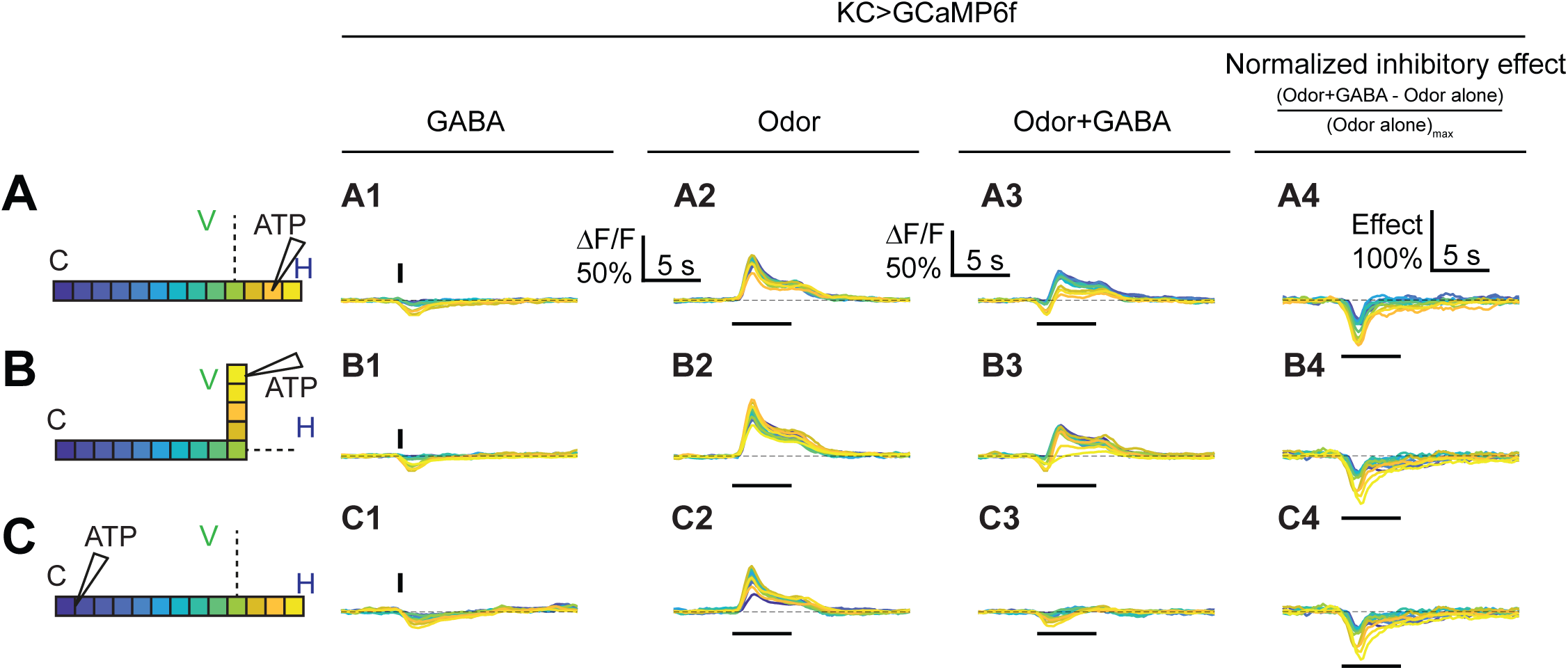
Time courses of GABA’s effect on KC activity. Rows: Local application of ATP (7.5 mM) in the horizontal lobe **(A1-A2)**, vertical lobe **(B1-B2)** or calyx **(C1-C2)**. Columns: Responses of KCs to GABA application **(A1-C1)**, the odor isoamyl acetate **(A2-C2)**, or both **(A3-C3). (A4-C4)** Normalized inhibitory effect of GABA application on KC responses to isoamyl acetate. Traces show the time course of the response in each segment of KCs, averaged across flies. Color-coded skeleton indicates which segments have traces shown (dotted lines mean the data is omitted for clarity; the omitted data appear Fig. 7). Vertical and horizontal bars indicate the timing of GABA and odor stimulation, respectively.

**Figure S4.**
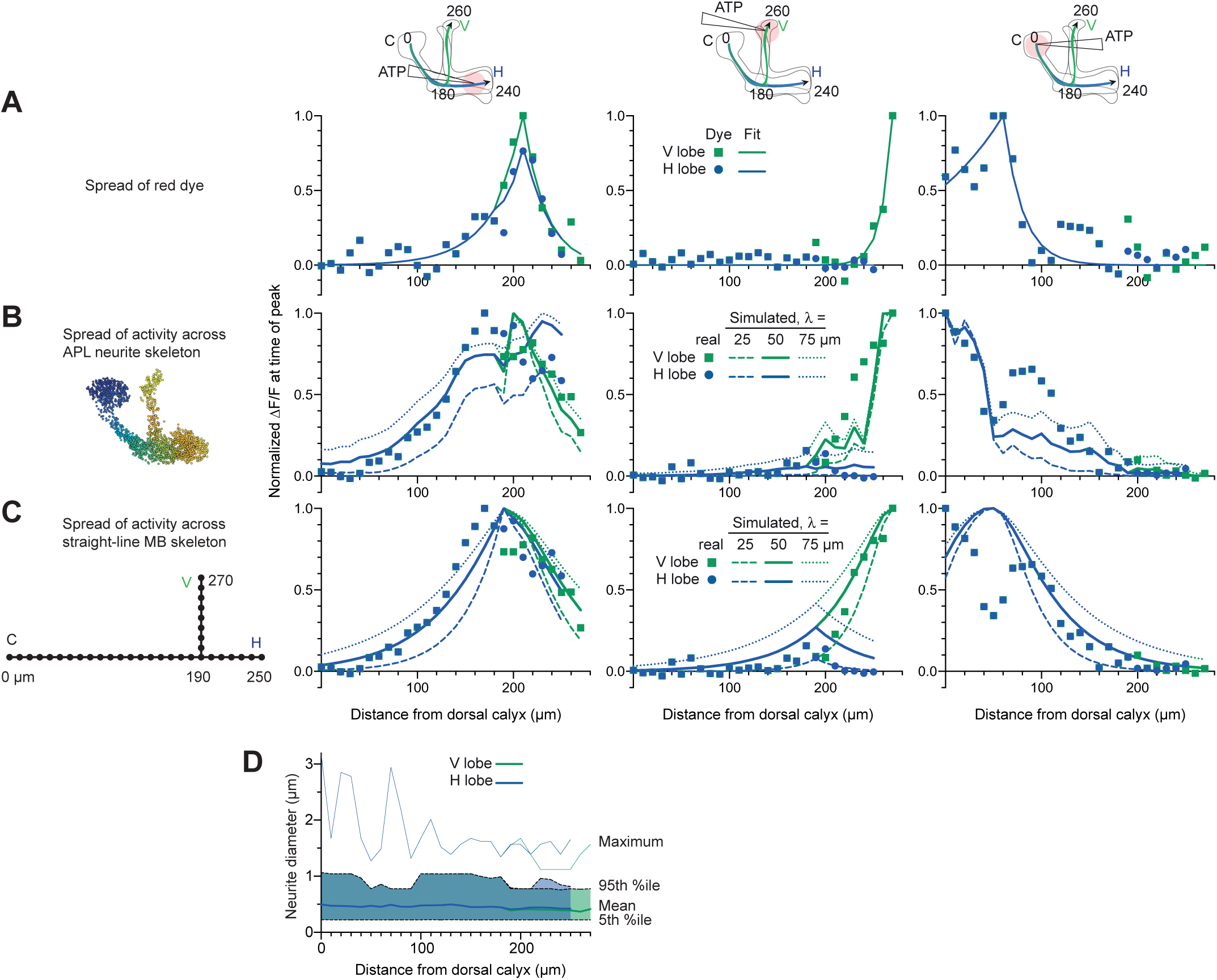
Additional data for Fig. 8. **(A-C)** Simulating local activation of APL with space constants 25, 50 and 75 µm. **(A)** and **(B)** repeated from **Fig. 8D,E** for comparison. **(C)** Simulated activity ignoring APL neurite morphology, treating the Voronoi cells as points on perpendicular straight lines. The same exponential decay applies as in B, except that the distance between points is the cityblock distance, i.e., the distance along the straight-line skeleton, not the APL’s neurite skeleton (as in **B**):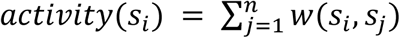. Note that 50 µm gives the best fit for horizontal lobe and calyx stimulation, while 25 µm gives the best fit for vertical lobe stimulation. **(D)** Diameter of neurites in each Voronoi cell of APL. Line shows mean, shaded area shows 5th to 95th percentile, and thin line above shows the maximum. Mean and percentiles were calculated by weighting each link in APL’s neurite skeleton by its length.

**Figure S5.**
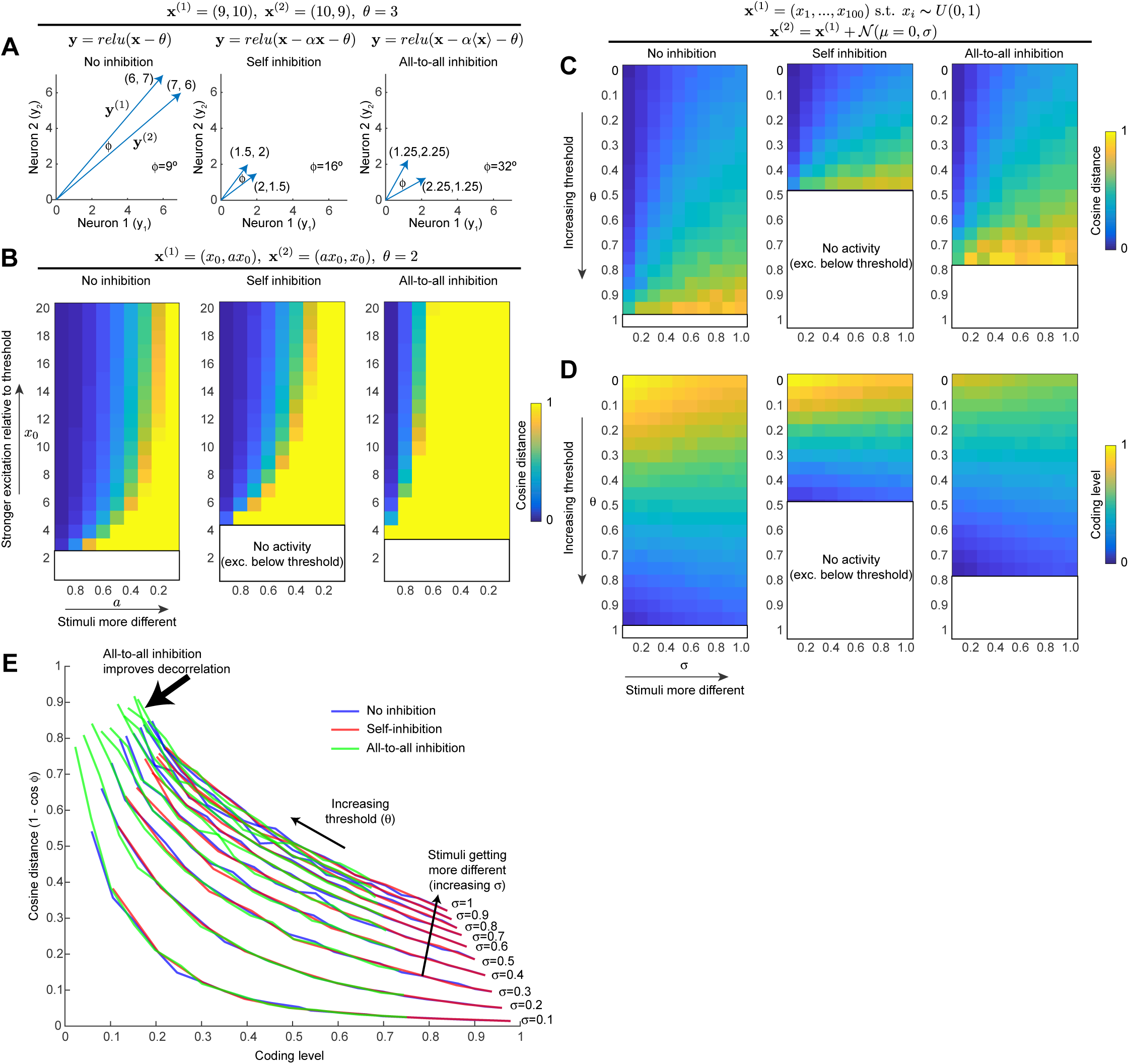
All-to-all inhibition decorrelates population activity better than self-inhibition (see Appendix for full description) **(A)** For a toy example of two neurons and two stimuli, self-inhibition improves the stimulus separation (i.e, increases angle ϕ between the two stimulus vectors) by bringing activity down closer to spiking threshold, but all-to-all inhibition improves the separation more. Note that to simplify the model, we model the inhibitory interneuron as being activated by the principal neurons’ dendrites, as occurs with Kenyon cells and APL. **(B)** Generalization of result in **(A)** to wider activity range in the two neurons. Heat maps show cosine distance for different values of *x*_0_ and *a* (increasing *x*_0_ means more excitation; decreasing a means the stimuli are more different). The self-inhibition panel is the same as the no-inhibition panel, just stretched vertically (i.e., self-inhibition acts as gain control), whereas the all-to-all inhibition panel expands the range of high cosine distances toward more similar stimulus pairs. Blank areas mean the activity is zero because excitation is below threshold, so cosine distance is undefined. **(C)** Generalization of **(A,B)** to population of 100 neurons. Stimulus 1 is sampled from a uniform distribution over (0, 1). Stimulus 2 is Stimulus 1 plus Gaussian noise (magnitude of noise governed by σ; higher σ means the two stimuli are more different). Heat maps show cosine distance for different thresholds (θ) and σ averaged over 100 trials. **(D)** Coding level (fraction of active neurons) for different thresholds and σ as in **(C)**; note how decreasing coding level (increased sparseness) correlates with higher cosine distance. **(E)** Each curve is one column (one value of σ) from the heat maps in **(C)** and **(D)**, showing how cosine distance improves with lower coding levels. Adding inhibition does not change this curve; it merely shifts the model to different points along the curve. All-to-all inhibition achieves higher cosine distance (more decorrelated activity) than self-inhibition or no inhibition.

## Supplementary Material

**Appendix:** Effect of self-inhibition vs. all-to-all inhibition on decorrelation of population activity

**Table S1:**
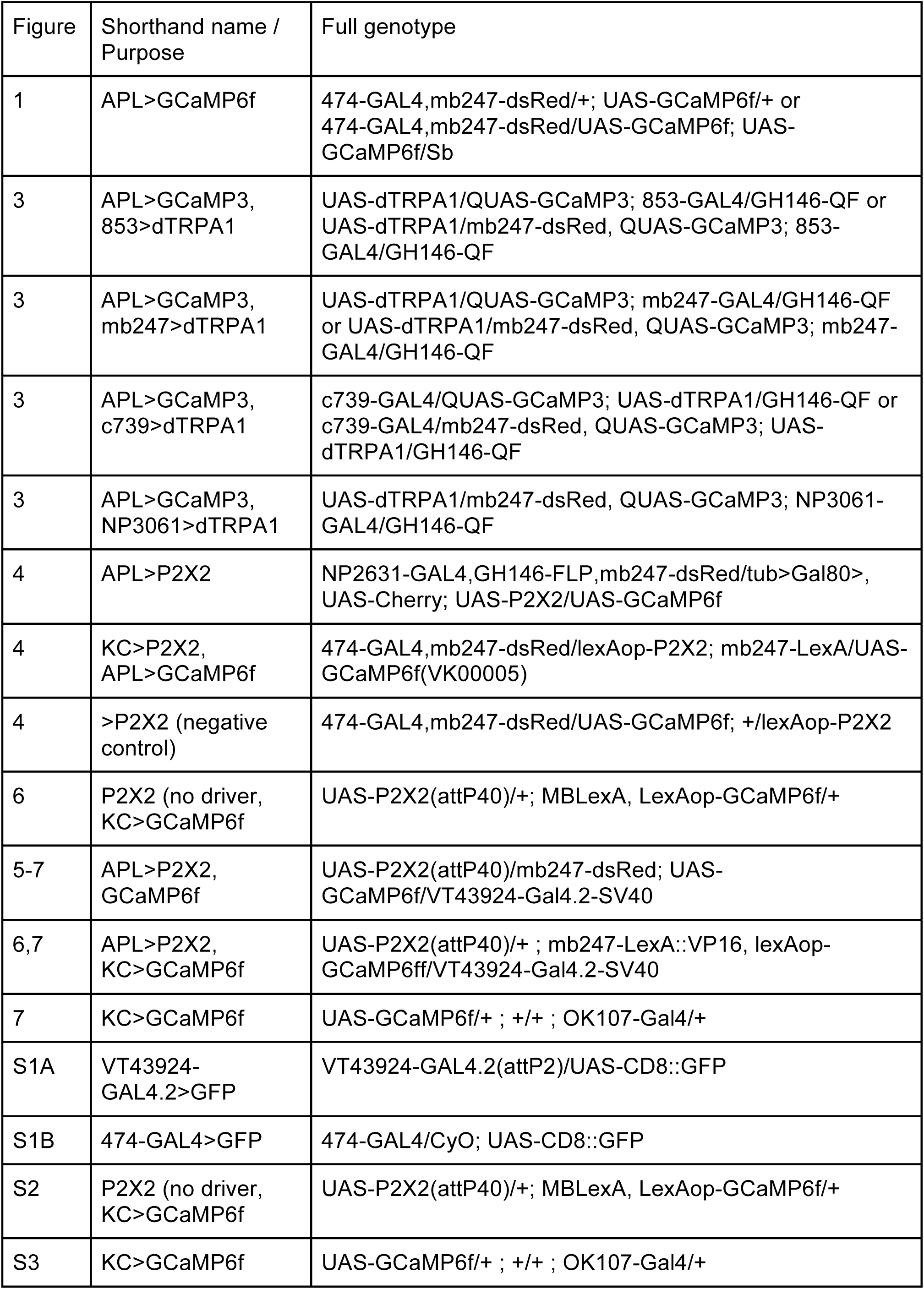
List of genotypes used

**Table S2.**
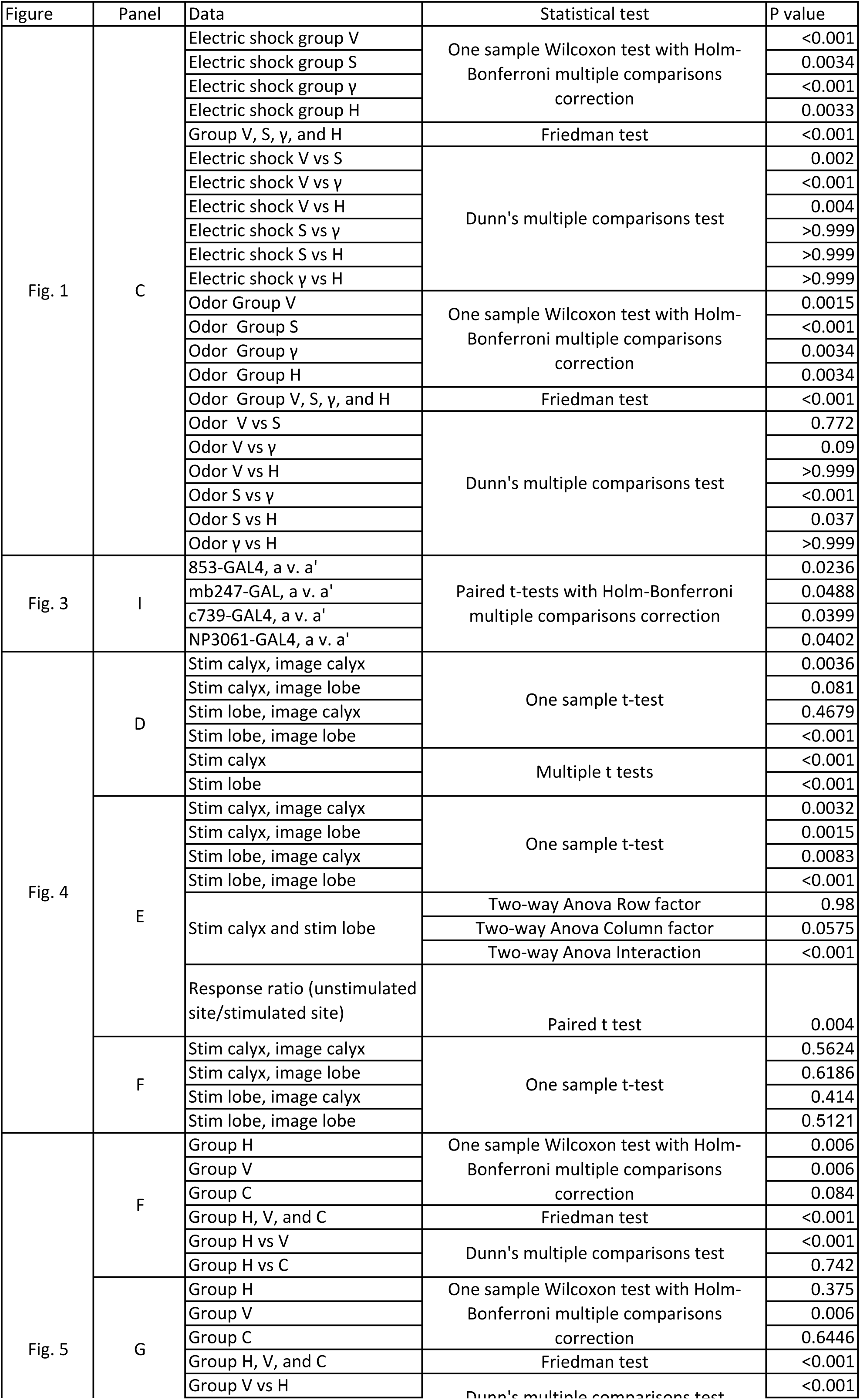
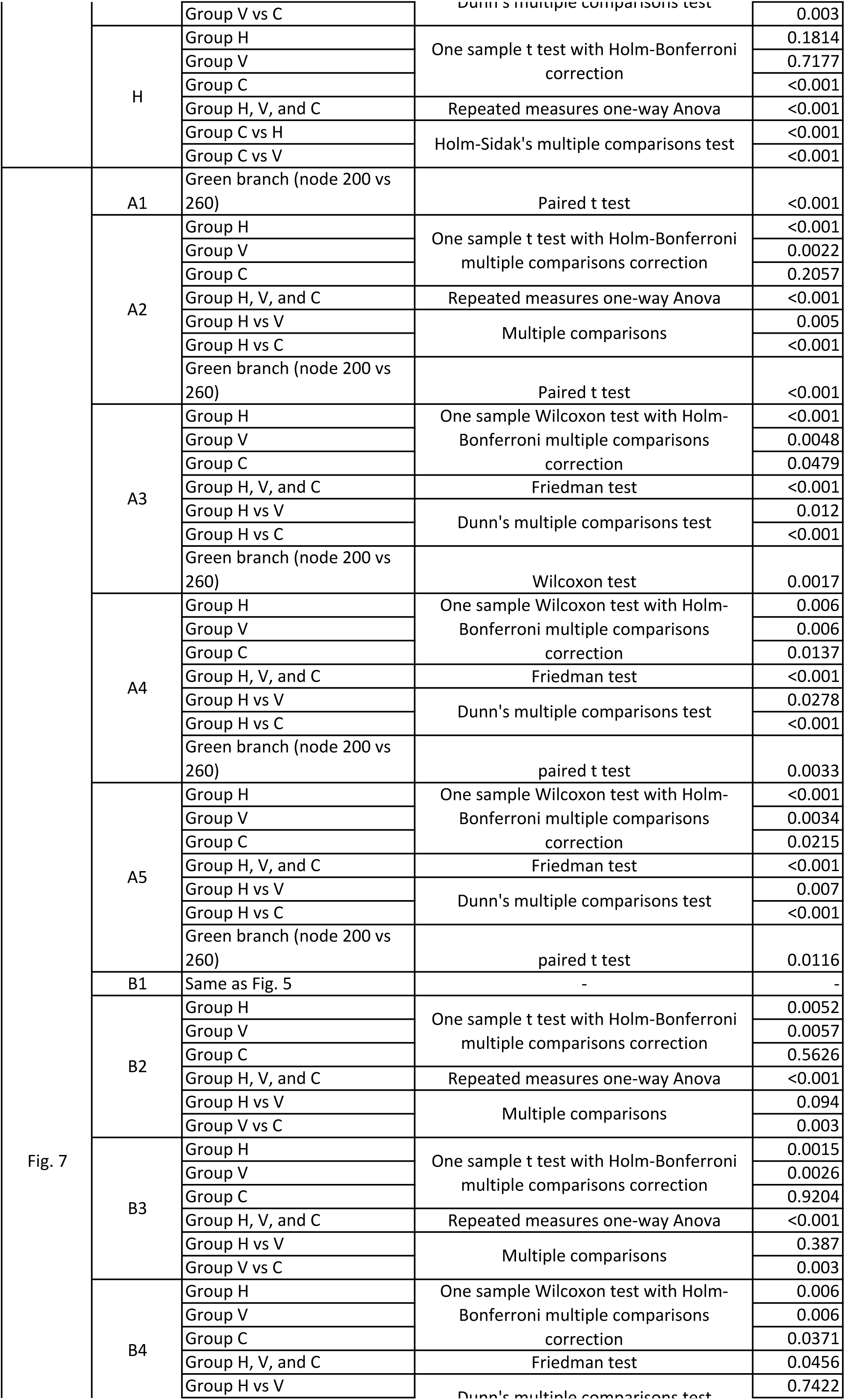
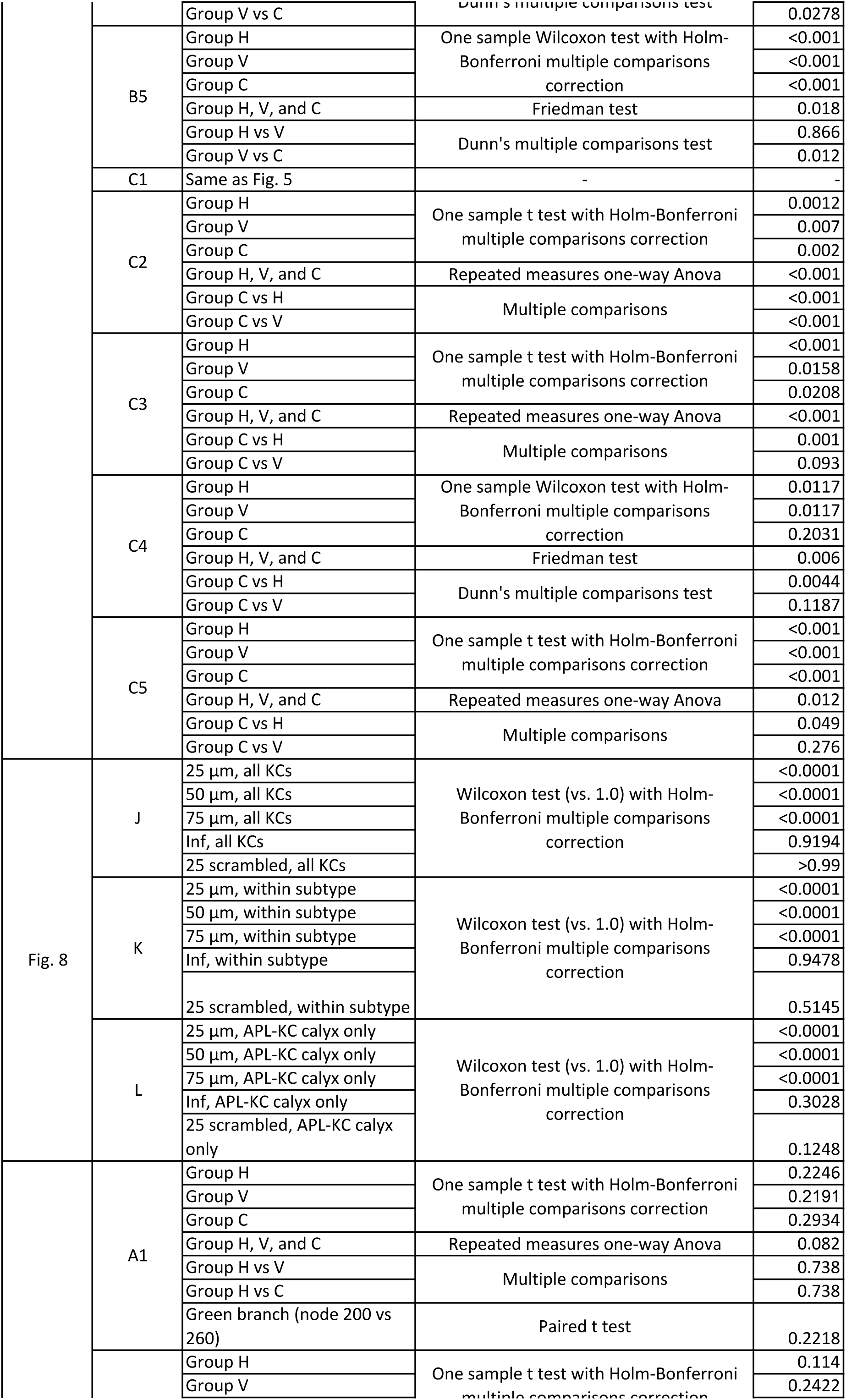
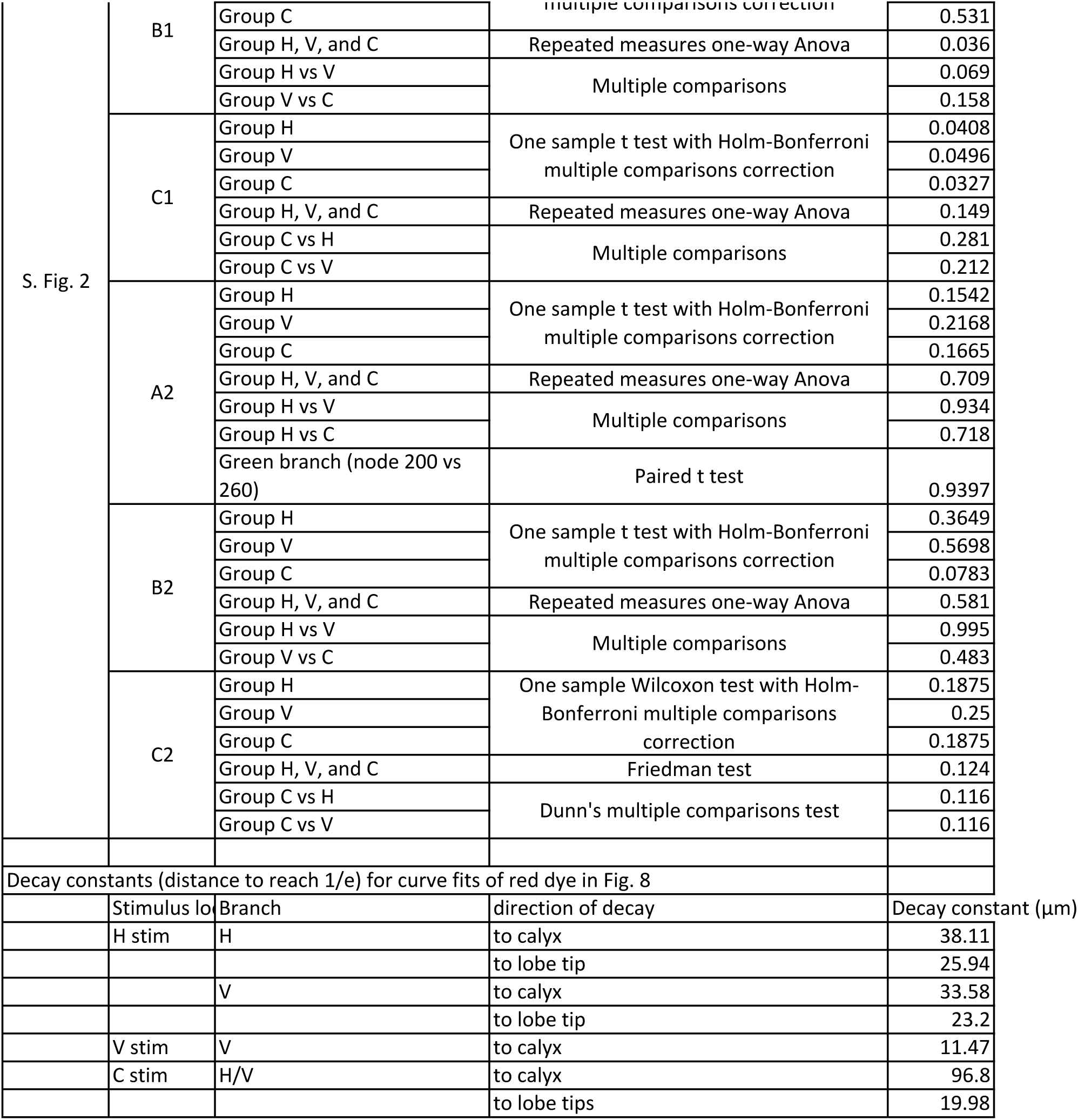
details of statistical analyses

## Appendix: Effect of self-inhibition vs. all-to-all inhibition on decorrelation of population activity

In this Appendix we develop and formalize our intuition for why all-to-all lateral inhibition is better than self-inhibition for decorrelating population activity, i.e., for enhancing contrast between stimuli.

Consider two linear rate-coding neurons such that the population response of the two neurons can be described as a vector **y** = (*y*_1_, *y*_2_), e.g., if neuron 1’s activity is 6 and neuron 2’s is 7, then population activity **y** = (6, 7). Take a simple case where **x** = (*x*_1_, *x*_2_) is the excitation and *θ* represents the spiking threshold:

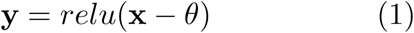

where relu is a rectified linear unit:

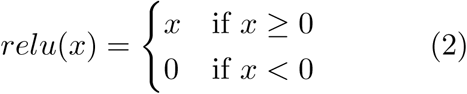

Suppose the two neurons respond to two similar stimuli, such that **x**^(1)^ = (9, 10) and **x**^(2)^ = (10, 9). If *θ* = 3, then **y**^(1)^ = (6, 7), **y**^(2)^ = (7, 6) (plotted on Fig. S5A).

Consider a form of inhibition in which the principal neurons excite the inhibitory neuron from their dendrites, not necessarily needing to depolarize above the spiking threshold. This is similar to the mushroom body, where Kenyon cells release acetylcholine from their dendrites and locally activate the inhibitory APL. If each principal neuron inhibits only itself by local inhibition - which we call self-inhibition - then

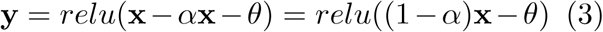

where *α* is the gain on the inhibitory neuron (0 *< α <* 1). Here, the self-inhibition is equivalent to decreasing the gain on the excitation. If we set *α* = 0.5 and *θ* = 3, then **y**^(1)^ = (1.5, 2) and **y**^(2)^ = (2, 1.5) (Fig. S5A). If in contrast, the inhibitory neuron adds up the activity of all principal neurons equally - which we call all-to-all inhibition - then

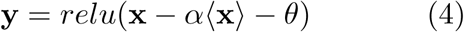

where ⟨**x**⟩ is the average excitation across the population. Using the parameters as for self-inhibition above, **y**^(1)^ = (1.25, 2.25) and **y**^(2)^ = (2.25, 1.25) (Fig. S5A).

We quantify the separation in the population response to these two stimuli using the cosine distance, i.e. 1 − cos(*ϕ*), where *ϕ* is the angle between the two vectors **y**^(1)^ and **y**^(2)^. It can be seen in Fig. S5A that self-inhibition increases the angle between the two vectors, but all-to-all inhibition increases it even more.

We generalize this example to a wider range of pairs of stimuli:

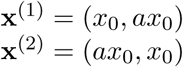

where *x*_0_ ranges from 1 to 20, *a* from 0.1 to 0.9. This generates pairs of stimuli that are very different when *a* is small, e.g., (1.5, 15) and (15, 1.5), or that are very similar when *a* is large, e.g. (13.5, 15) and (15, 13.5). We set *θ* = 2.

In the no-inhibition case, cosine distances are higher at lower *x*_0_ because this brings activity closer to the threshold (Fig. S5B). That is, (3, 4) and (4, 3) become more different from each other once a threshold of 2 is applied, turning the vectors to (1, 2) and (2, 1), but the effect is not so strong for turning (13, 14) and (14, 13) into (11, 12) and (12, 11).

Compared to the no-inhibition case, self-inhibition increases cosine distances for a given *x*_0_ and *a*. However, this is functionally equivalent to simply raising *θ* dynamically for higher *x*_0_. Visually, this is seen on Fig. S5B by the way the self-inhibition panel is simply the no-inhibition panel stretched vertically. That is, self-inhibition decorrelates mainly by using gain control to bring down the activity closer to threshold.

In contrast, all-to-all inhibition improves decorrelation for higher *a* (i.e., more similar stimuli). Visually, this is seen on Fig. S5 as the zone of large distances expanding to the right. That is, all-to-all inhibition can make similar stimuli appear more distinct.

Next we generalize these intuitions to larger populations of neurons. Consider a population of 100 neurons that receive excitation **x**:

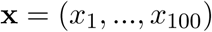

Stimulus 1 consists of random numbers between 0 and 1 (uniform distribution):

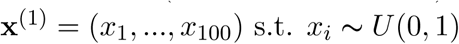

Stimulus 2 is similar to Stimulus 1, which we simulate by making it equal to Stimulus 1 plus Gaussian noise with mean 0 and standard deviation *σ*:

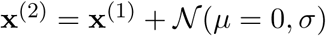

We apply these excitations to the no-inhibition, self-inhibition, or all-to-all inhibition conditions, with *θ* ranging from 0 to 0.1 and *σ* ranging from 0.1 to 1. For all three conditions, cosine distances increase for higher *σ* because the two stimuli are more different (Fig. S5C). They also increase for higher *θ* because more neurons are silenced (i.e., the coding level, or fraction of responsive neurons, is reduced) (Fig. S5D).

Compared to the no-inhibition condition, self-inhibition reaches similar cosine distances but at lower *θ*, consistent with our earlier conclusion that self-inhibition mainly functions equivalently to dynamically adjusting the threshold. In other words, self-inhibition helps decorrelate by pushing neurons’ activity down closer to, or indeed below, the threshold. In contrast, all-to-all inhibition (1) reaches higher cosine distances than the other two conditions and (2) allows higher cosine distances at lower thresholds.

This can be more clearly understood by plotting cosine distance against coding level for different values of *σ*. In Fig. S5E, each line is one column (one value of *σ*) in the heat maps in Fig. S5C,D. Within this range of parameters, the fundamental relation between coding level and cosine distance is fixed for a given *σ*. That is, for a given *σ*, all three conditions follow the same curve on the cosine distance vs. coding level graph. What differs is *where* on the curve they fall. The decorrelating benefit created by all-to-all inhibition arises in part from pushing the population activity up and to the left along the curve. A second benefit arises at the upper left end of the curve: this is where population activity falls silent because *θ* is so high (corresponding to the blank rows in the heat maps). All-to-all inhibition allows non-zero population activity to persist a bit further on this curve, where the other two conditions have already fallen silent. In contrast, self-inhibition merely moves population activity further along the curve at lower values of *θ* without moving beyond the range allowed without inhibition.

